# Modeling Genetic Epileptic Encephalopathies using Brain Organoids

**DOI:** 10.1101/2020.08.23.263236

**Authors:** Daniel J. Steinberg, Afifa Saleem, Srinivasa Rao Repudi, Ehud Banne, Muhammad Mahajnah, Jacob H. Hanna, Peter L. Carlen, Rami I. Aqeilan

## Abstract

Epileptic encephalopathies (EEs) are a group of disorders associated with intractable seizures, brain development and functional abnormalities, and in some cases, premature death. Pathogenic human germline biallelic mutations in tumor suppressor WW domain-containing oxidoreductase (WWOX) are associated with a relatively mild autosomal-recessive spinocerebellar ataxia-12 (SCAR12) and a more severe early infantile WWOX-related epileptic encephalopathy (WOREE). In this study, we generated an *in-vitro* model for EEs, using the devastating WOREE syndrome as a prototype, by establishing brain organoids from CRISPR-engineered human ES cells and from patient-derived iPSCs. Using these models, we discovered dramatic cellular and molecular CNS abnormalities, including neural population changes, cortical differentiation malfunctions, and Wnt-pathway and DNA-damage response impairment. Furthermore, we provide a proof-of-concept that ectopic WWOX expression could potentially rescue these phenotypes. Our findings underscore the utility of modeling childhood epileptic encephalopathies using brain organoids and their use as a unique platform to test possible therapeutic intervention strategies.

## Introduction

Epilepsy is a neurological disorder characterized by a chronic predisposition for the development of recurrent seizures (Fisher *et al*., 2014; Aaberg *et al*., 2017). Epilepsy affects around 50 million people worldwide and is considered the most frequent chronic neurologic condition in children (Aaberg *et al*., 2017; Blumcke *et al*., 2017). Approximately 40% of seizures in the early years of life are accounted for by early infantile epileptic encephalopathies (EIEEs), which are pathologies of the developing brain, characterized by intractable epileptiform activity and impaired cerebral and cognitive functions (Lado *et al*., 2013; Shao and Stafstrom, 2016; Nashabat *et al*., 2019). Several genes have been implicated in causing EIEEs (McTague *et al*., 2016). In recent years, autosomal-recessive mutations in the WW domain-containing oxidoreductase *(WWOX)* gene, are increasingly recognized for their role in the pathogenesis of EIEEs (Piard *et al*., 2018; Nashabat *et al*., 2019). WWOX, a tumor suppressor that spans the chromosomal fragile site FRA16D, is highly expressed in the brain, suggesting an important role in central nervous system (CNS) biology (Abu-Remaileh *et al*., 2015). In 2014, WWOX was implicated in the autosomal-recessive spinocerebellar ataxia-12 (*SCAR12*) (Gribaa *et al*., 2007; Mallaret *et al*., 2014), and in the WWOX-Related Epileptic Encephalopathy (WOREE Syndrome; also termed EIEE28) (Abdel-Salam *et al*., 2014; Ben-Salem *et al*., 2015; Mignot *et al*., 2015). Both disorders are associated with a wide variety of neurological symptoms, including seizures, intellectual disability, growth retardation and spasticity, but differ in severity, onset and by underlying types of mutations. The WOREE syndrome is considered more aggressive, appearing as early as 1.5 months and associating with more extreme genetic changes (Piard et al., 2018). This observation may imply that both syndromes can be considered as a continuum. Alongside seizures, WOREE-patients may present with global developmental delay, progressive microcephaly, atrophy of specific CNS components and premature death.

Although modeling WWOX loss of function in rodents has shed some lights on the roles of WWOX in brain function (Aqeilan *et al*., 2007, 2008; Suzuki *et al*., 2009; Mallaret *et al*., 2014; Tanna and Aqeilan, 2018; Tochigi *et al*., 2019), the genetic background and brain development of a specific patient cannot be modeled in a mouse, but is inherent and retained in patient-derived induced pluripotent stem cells (iPSCs). In an effort to bypass the comprehensible lack of availability of EE brain samples, including those of WOREE-patients’, we utilized genome editing and reprogramming technologies to recapitulate the genetic changes seen in WOREE and SCAR12 patients in human PSCs. We then generated brain organoids, 3D neuronal cultures, that recapitulate much of the brain’s spatial organization and cell type formation, and have neuronal functionality *in-vitro* (Amin and Paşca, 2018; Sidhaye and Knoblich, 2020). This allowed us to model features of the development and maturation of the CNS and its complex circuitry, in a system that is more representative of the *in-vivo* human physiology than 2D cell cultures. Using this platform, we identified severe defects in neural cell populations, cortical formation, and electrical activity, and tested possible rescue strategies. This approach has resulted in a deeper understanding of WWOX physiology and pathophysiology in the CNS, laying the foundation for developing more appropriate treatments, and supports the concept of using human brain organoids for modeling other human epileptic diseases.

## Results

### Generation and characterization of WWOX knockout cerebral organoids

To shed light on the pathogenesis of EIEE, we studied the WOREE syndrome as a prototype model using brain organoids. The role of WWOX in the development of the human brain in a controlled genetic background was investigated by generating WWOX knockout (KO) clones of the WiBR3 hESC line using the CRISPR/Cas9 system (Abdeen *et al*., 2018). Immunoblot analysis was used to assess WWOX expression in these lines (Supplementary figure S1A). Two clones that showed consistent undetectable protein levels of WWOX throughout our validations were picked for the continuation of the study (Supplementary figure S1A and S1B) – WWOX-KO line 1B (WKO-1B, from here on KO1) and WKO-A2 (from here on KO2). Sanger sequencing confirmed editing of WWOX sequence at exon 1 (Supplementary figure S1C). Furthermore, to confirm cell-autonomous function of WWOX, we restored *WWOX* cDNA into the endogenous *AAVS* locus of WWOX-KO1 hESC line and examined reversibility of the phenotypes (see Methods section and Supplementary figure S1D). The KO1-AAV4 line was selected for generating COs for having a strong and stable expression of WWOX throughout our validations (data not shown), and from here on is called W-AAV. These lines were practically indistinguishable from the parental cell line (WiBR3 WT) in terms of morphology and proliferation throughout the culture period (data not shown).

To investigate how depletion of WWOX affects cerebral development in a 3D context, we differentiated our hESCs into cerebral organoids (COs), using an established protocol (Lancaster *et al*., 2013; Lancaster and Knoblich, 2014). COs from all genotypes showed comparable gross morphology and development at all stages (data not shown). Next, we investigated the expression pattern of WWOX in the developing brain at different time points by co-staining with markers of the two major populations found in the organoids – neuronal progenitor cells and neurons. As seen in week 10, WWOX expression was specifically localized to innermost layer of the ventricular-like zone (VZ), which is composed of SOX2^+^ cells, corresponding to radial glia cells (RGs), the progenitors of the brain, and not in the surrounding cells (Figure 1A). This finding is in concordance with previous work showing limited WWOX expression during early steps of mouse cortical development (Chen *et al*., 2004). Furthermore, even in later time points, such as week 24, when the VZ structure is lost, WWOX expression is found mainly in SOX2^+^ cells (Figure 1B). Importantly, WWOX expression was not seen in COs generated from the WWOX-KO lines, although similar levels of expression of the other markers such as SOX2 and Neuron-specific class III β-Tubulin (TUBB3 or TUJ1) were observed (Figures 1A and 1B, Supplementary figure S1B and S1E). Interestingly, the W-AAV COs, in which WWOX expression is driven by human ubiquitin promotor (UBP), exhibited high WWOX levels in the VZ, as expected, though other cellular populations also showed prominent WWOX expression (Supplementary figure S1E).

**Figure 1.**
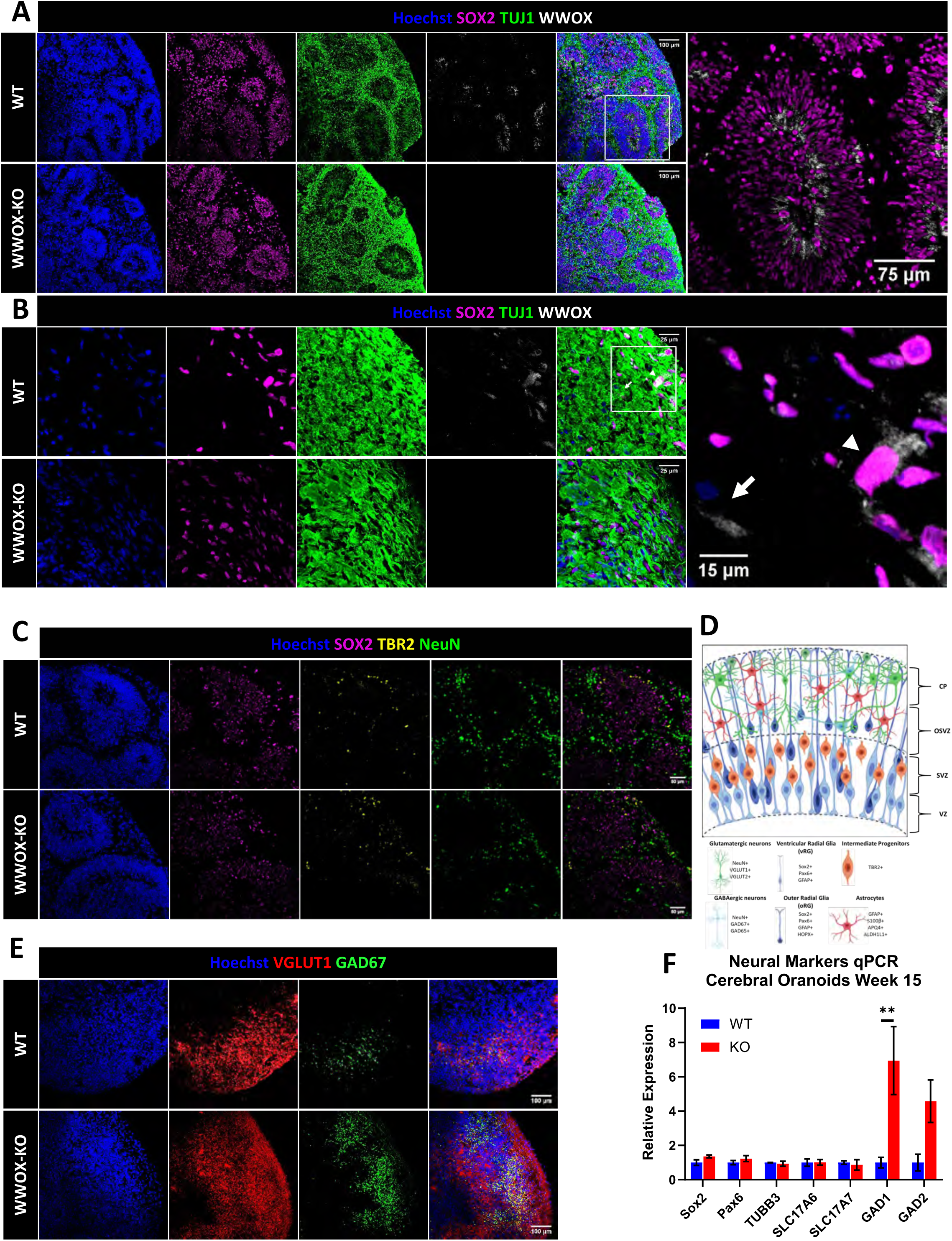
Generation and characterization of WWOX Knock-out Cerebral Organoids. A) Week 10 cerebral organoids (COs) stained for the progenitor marker Sox2, neuronal marker Tuj1 and WWOX. B) Week 24 COs stained for SOX2, TUJ11 and WWOX. Arrowhead denotes a SOX2^+^ cell that express WWOX, arrow denotes Sox2^-^ cell that expresses WWOX. C) Week 10 COs stained for the mature neural marker NeuN, intermediate neurons marker TBR2 (EOMES) and progenitor marker SOX2 *(WT: n=4, KO: n=3)*. D) Schematic representation of the different population forming the VZ and the adjacent surrounding. E) Immunofluorescent (IF) staining for the glutamatergic neurons marker VGLUT1 and GABAergic neurons marker GAD67 (GAD1) *(WT: n=4, KO: n=3)*. F) qPCR analysis for the assessment of expression levels of different neural markers in 15 weeks COs: SOX2 and PAX6 (progenitor cells), SLC17A6 and SLC17A (VGLUT2 and VGLUT1; glutamatergic neurons) and GAD1 and GAD2 (GAD67 and GAD65; GABAergic neurons). Y-axis indicated relative expression fold change. Data are represented as mean ± SEM.

Next, we further examined the development of cerebral structures. In WT organoids, the VZ, which is composed of SOX2^+^ cells, is surrounded by intermediate progenitors (IP; TBR2^+^ cells, also known as EOEMS), marking the presence of the subventricular zone (SVZ). Outside this layer is the cortical plate (CP), composed mainly by neurons (NeuN^+^ cells) (Figure 1D). In week 10 COs, no visible differences in the composition or formation of the VZ and the surrounding structures were observed (Figure 1C), suggesting similar proportions of these populations. This was also supported by measuring RNA expression levels of the progenitor markers *SOX2* and *PAX6*, and the neuronal marker *TUBB3* (Figure 1F and Supplementary figure S1F). This surprising observation led us to further examine the two major neuronal sub-populations found in COs – Glutamatergic (marked by vesicular glutamate transporter 1; VGLUT1) and GABAergic neurons (marked by glutamic acid decarboxylase 67; GAD67). Immunostaining revealed that although VGLUT1 expression remained similar, a marked increase in expression of GAD67 was observed in KO COs compared to WT (Figure 1E). In contrast, WWOX restoration (W-AAV) significantly reversed this imbalance (Supplementary figure S1G). RNA levels of *SLC17A6 (VGLUT2), SLC17A7 (VGLUT1), GAD1 (GAD67) and GAD2 (GAD65)* followed the same direction (Figure 1F, Supplementary figure S1F).

These findings suggest that during human embryonic development, WWOX expression is limited to the cells of the basal layer of the VZ and that WWOX depletion does not affect the VZ-SVZ-CP architecture, but did disrupt the balance between glutamatergic and GABAergic neurons.

### WWOX-depleted cerebral organoids exhibited hyper-excitability and epileptiform activity

Cerebral organoids give rise to neurons that have previously shown electrophysiological functionality (Trujillo *et al*., 2019). To characterize the functional properties of the WWOX-KO COs, we performed local field potential recordings (LFP) in 7-week COs slices. Electrodes were positioned 150µm away from the edge of the slice (Supplementary figure S2A) to avoid areas potentially damaged by slice preparation. Sample traces of the WT and KO COs revealed visible differences between the two lines under baseline conditions (Figure 2A, left). The mean spectral power of field recordings further showed an overall increase in power of the KO COs in the 0.25-1 Hz low frequency range (Figure 2B) and a decrease in the 30-79.9 high frequency range (Supplementary figures S2B and S2C). The oscillatory power (OP) was quantified by the area under the curve, which was significantly higher than the WT line, under baseline conditions (Figure 2C). Over time, the OP of the KO lines decreased significantly while the WT line’s OP stayed the same (Supplementary figure S2E), suggesting a developmental delay in the KO line.

**Figure 2.**
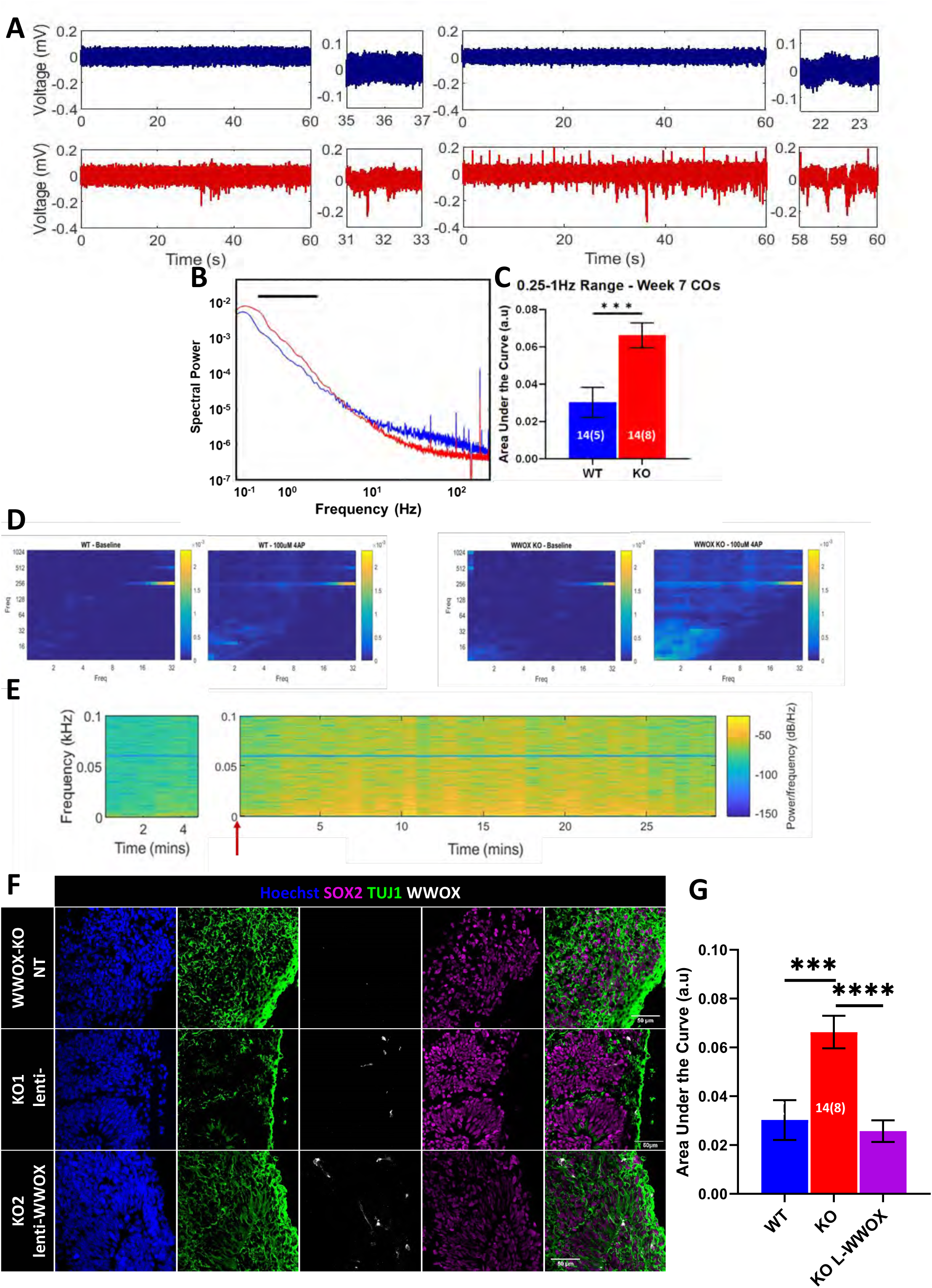
WWOX-KO Cerebral Organoids Demonstrated hyper-excitability and epileptiform activity. Sample recordings from 7-week old hESC-derived Cerebral Organoids (COs). (A) Traces suggest that WWOX-KO COs show increased activity compared to baseline in both groups. (B) Mean spectral power of wildtype (WT) and 2 knockout lines at week 7 in baseline conditions. (C) Normalized area under the curve of the mean spectral power in (A) for the 0.25-1 Hz frequency range. Data represented by mean ± SEM. The two-tailed unpaired Student’s t-test was used to test statistical significance. The numerals in all bars indicate the number of analyzed slices and organoids (i.e. slices (organoids)). (D) CFC analysis shows increased coupling in the δ: HFO frequency pairs. (E) Sample spectrogram of WWOX KO slice shows a gradual increase in activity upon addition of 4AP (marked by red arrow) for up to 2 min, and a decrease after 25 min. All traces were filtered with a 60 Hz notch filter and 0.5 Hz high-pass filter. F,G WWOX’s coding sequence was re-introduced into week 6 WWOX-KO COs using lentiviral transduction (lenti-WWOX). (F) Immunofluorescent staining showing WWOX expression in different populations in WWOX-KO organoids following infection with lentivirus. (G)Normalized area under the curve of the mean spectral power of WT line, 2 KO lines and 2 KO lines infected with lenti-WWOX at week 7 in baseline condition, for the 0.25-1 Hz frequency range. Data represented by mean ± SEM. The numerals in all bars indicate the number of analyzed slices and organoids (i.e. slices (organoids)).

To further measure the hyper-excitability of the KO line, 100µM 4-AP, a commonly used convulsant for seizure induction, was applied to the slices during recordings. While 4-AP did show changes in LFP recordings for both WT and KO lines (Figure 2A, right), the KO line showed significantly increased activity, which was otherwise absent in WT traces (Supplementary figure S2D). The effect of 4-AP on spectral power became evident 5 minutes after its addition, as indicated by the sample spectrogram (Figure 2E). Cross-frequency coupling of the sample traces for both WT and KO lines in the presence of 4- AP revealed an increase the δ: HFO frequency pairs – an attribute which has previously been used to characterize and classify seizure sub-states (Figure 2D) (Guirgis *et al*., 2013). Importantly, transduction of the lentivirus containing WWOX resulted in a recovery of the KO line, with respect to the mean power spectral density (Figure 2F-G).

### WWOX-depleted cerebral organoids exhibited impaired astrogenesis and DNA-damage response

It is widely accepted that an imbalance between excitatory and inhibitory activity in the brain is a leading mechanism for seizures, but this does not necessarily mean neurons are the only population involved. It is well known that brain samples from epileptic patients show signs of inflammation, astrocytic activation and gliosis (Cohen-Gadol *et al*., 2004; Thom, 2009), which can be a sole histopathological finding in some instances (Blumcke *et al*., 2017). Whether this phenomenon is a results of the acute insult or a cause of the seizures, is still debatable (Vezzani *et al*., 2011; Robel *et al*., 2015; Rossini *et al*., 2017; Patel *et al*., 2019). Furthermore, recent work has demonstrated astrogliosis in the brain of *Wwox*-null mice (Hussain *et al*., 2019).

To address this, we used immunofluorescence staining to visualize the astrocytic markers glial fibrillary acidic protein (GFAP) and S100 calcium-binding protein B (S100β) in week 15 and week 24 COs (Figure 3A and Supplementary figure S1A). This revealed a marked increase in astrocytic cells in WWOX-KO COs, that progressed through time, and was partially reversed in W-AAV COs (Figure 3B and Supplementary figure S1B). It is notable that GFAP also marks RGs (Middeldorp *et al*., 2010), which are abundant in week 15, but are reduced in number in week 24, which could be a source of noise at early stages.

**Figure 3.**
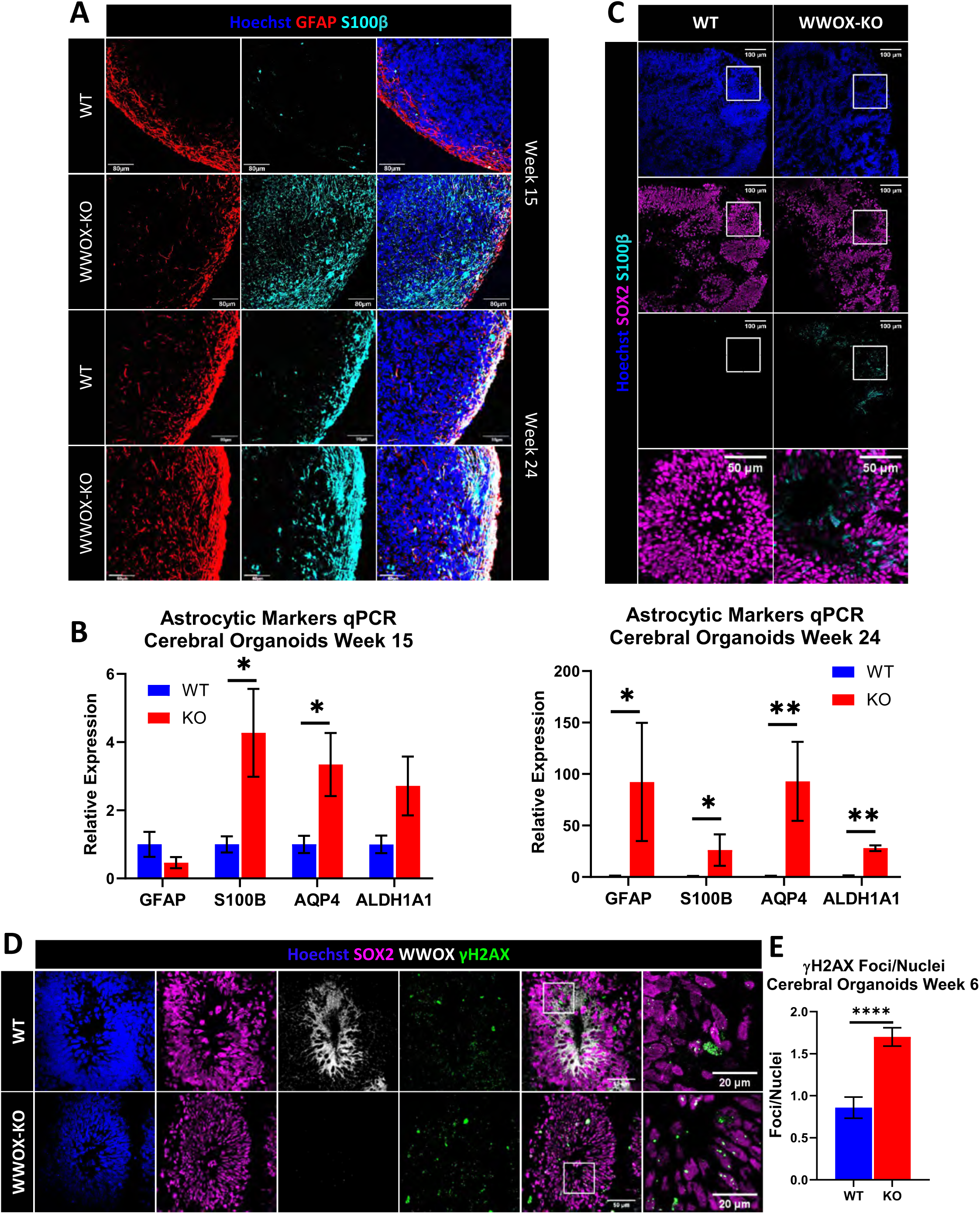
WWOX-KO Cerebral Organoids Showed Impaired Astrogenesis and DNA-damage Response. A) Week 15 and week 24 COs stained for the astrocytic and radial glia marker GFAP, and the astrocyte-specific marker S100β *(WT W15: n=3, KO 15: n=4. WT W24: n=4, KO W24: n=2)*. B) qPCR analysis of astrocytic markers in COs at week 15 (top) and week 24 (bottom). Y-axis indicated relative expression fold change. Data are represented as mean ± SEM. (*WT W15: n=4, KO 15: n=4. WT W24: n=4 from 2 individual batches, KO W24: n=3.)*. C) IF staining of week 6 COs for astrocytic markers in the surrounding of VZs (WT: n=6 organoids from 2 individual batches, KO: n=6 organoids from 2 individual batches). D) Staining for the DNA damage marker γH2AX in week 6 in the nuclei of cells in the VZ at physiological conditions *(WT: n=8 organoids from 3 individual batches, KO: n=12 organoids from 3 individual batches)*. E) Quantification of γH2AX foci in the nuclei of cells composing the innermost layer of the VZ, normalized to the total number of nuclei in this layer. Data are represented as mean ± SEM *(WT: n=8* organoids from 3 individual batches, KO: n=12 organoids from 3 individual batches).

Astrocytes arise from two distinct populations of cells in the brain: the RG cells, switching from neurogenesis to astrogenesis, or from astrocytes progenitor cells (APCs) (Zhang *et al*., 2016; Blair, Hockemeyer and Bateup, 2018). To track back these differences in astrocyte markers, we compared 6- and 10-week old COs. We found that in week 6 COs, where no astrocytic markers were detected in WT organoids, a significant expression of S100β was observed in the basal layer of VZ region (Figure 3C). At week 10, we observed expression of S100β in both WT and KO organoids, but when co-staining with the cell proliferation marker Ki67, we did not detect a significant difference in double-positive cells suggesting similar glial proliferation (Supplementary figure S3C). Quantification of Ki67^+^ nuclei, together with SOX2^+^ nuclei revealed that although the proportions of SOX2 remained intact in WT compared to WWOX-KO (18.5% in WT, 95% CI=14.2-22.81; 19.5% in KO, 95% CI=15.29-22.81), the proportions of proliferating cells (9.5% in WT, 95% CI=6.63-12.35; 4.9% in KO, 95% CI=2.76-7.13) and Ki67^+^/SOX2^+^ double-positive cells were diminished (51.83% in WT, 95% CI=41.18-62.48; 27.09% in KO, 95% CI=14.1-40.1) (Supplementary figure S3D). These findings imply that the initial increase in astrocytic markers in WWOX-depleted COs is likely due to enhanced differentiating RGs rather than proliferating APCs.

This peculiar behavior of the RGs in KO COs led us to take a closer look at their functionality by examining their physiological DNA damage response (DDR), a signaling pathway in which WWOX is known to be directly involved (M. Abu-Odeh, Salah, *et al*., 2014; Abu-odeh, Hereema and Aqeilan, 2016). To this end, we stained for γH2AX, a surrogate marker for DNA double strand breaks. We found a marked accumulation of γH2AX foci in the nuclei of SOX2^+^ cells in the innermost layer of the VZ, averaging 1.7 foci/nuclei [*95% CI=1.48-1.92]* in WWOX-KO, compared to 0.86 foci/nuclei [*95% CI=0.6-1.12*] in the age-matched WT COs (Figures 3D and 3E). To further verify our observations, we stained for p53-binding protein 1 (53BP1), another DDR signaling marker, and found increased foci in WWOX-depleted organoids (Supplementary figure S3E). These findings are consistent with WWOX direct role in DDR signaling (Aqeilan, Abu-Remaileh and Abu-Odeh, 2014; Hazan, Hofmann and Aqeilan, 2016). Importantly, W-AAV COs presented with improved DDR (Supplementary figure S3F). Intriguingly, a higher number of diffuse nuclear γH2AX staining was observed, suggestive of increased apoptosis (Rogakou *et al*., 2000; Solier and Pommier, 2009), a phenomenon previously reported upon WWOX overexpression (Chang *et al*., 2001, 2005; Chang, Doherty and Ensign, 2003; Aqeilan *et al*., 2004; Del Mare, Salah and Aqeilan, 2009). The γH2AX foci were found in highly proliferating cells in the VZ, which was observed by co-staining with Ki67 - 18.6% of SOX2^+^ cells were double positive in KO COs [95% CI=15-22%] compared to 11.9% [95% CI=8-15%] (Supplementary figure S3G, 3H).

In conclusion, WWOX-KO COs present with a progressive increase in astrocytic number, likely due to enhanced differentiating RGs, and with increased DNA damage in neural progenitor cells.

### RNA-sequencing of WWOX depleted cerebral organoids revealed major differentiation defects

In effort to examine the molecular profiles, we performed whole-transcriptome RNA sequencing (RNA-seq) analysis on week 15 WT and KO COs. Albeit the known heterogeneity of brain organoids, Principal Component Analysis (PCA) separated the sample into two distinct clusters (Supplementary figure S4A). The analysis revealed 15,370 differentially expressed genes, of which 1,246 genes were upregulated in WWOX-KO COs, and showed a greater than 1.2 fold change (*FC>1.2)* and a significant p-value (*P-Value<0.01)*, and 1,021 genes were down-regulated (*FC<1/1.2, P-Value<0.01)* (Supplementary figure S4B; Supplementary Tables 1-2). Among the top 100 upregulated genes, we found genes related to neural populations such as GABAergic neurons (*GAD1, GRM7, LHX5*) and astrocytes (*AGT, S100A1, GJA1, OTX2*), and to neuronal processes like calcium signaling (*HRC, GRIN2A, ERBB3, P2RX3, HTR2C, PDGFRA*) and axon guidance (*GATA3, DRGX, ATOH1, NTN1, SHH, RELN, OTX2, SLIT3,GBX2, LHX5*) (Figure 4A). In the top 100 downregulated genes were genes related to GABA receptors (*GABRB3, GABRB2*), autophagy *(IFI16, MDM2, RB1, PLAT, RB1CC1)* and the mTOR pathway (*EIF4EBP1, PIK3CA, RB1CC1*) (Figure 4A).

**Figure 4.**
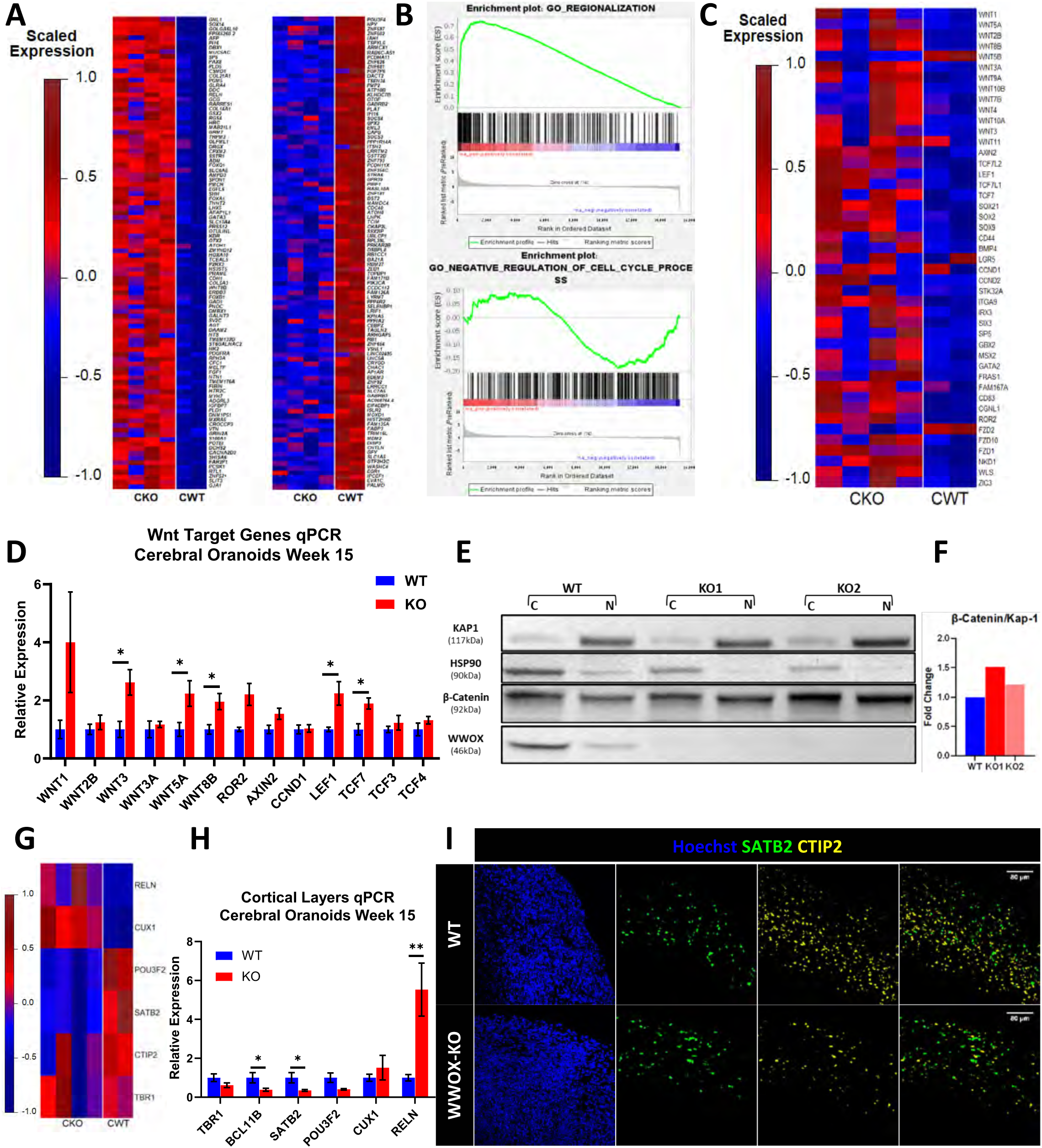
Cerebral Organoids RNA-sequencing Revealed Major Differentiation Defects. RNA sequencing (RNA-seq) of week 15 COs and transcriptome analysis *(WT: n=2, KO: n=4)*. A) Heatmap of 100 upregulated genes (left panel) and 100 downregulated genes (right panel) selected by highest fold change. B) Gene-set enrichment analysis (GSEA) revealed enrichment of genes related to regionalization of the organoids, and decreased expression of genes related to negative regulation of the cell cycle. C) Heatmap of Wnt-pathway related genes in 15-weeks COs. D) qPCR analysis for selected Wnt genes validating the results of the RNA-seq. Y-axis indicated relative expression fold change. Data are represented as mean ± SEM (*WT W15: n=4, KO 15: n=4.)* E) Week 16 COs were sub-fractionated into a cytoplasmic (C) and nuclear (N) fractions. The experiment was run twice with a total of 2 WT organoids and 4 KOs organoids (2 for each KO line). KAP-1 marks the nucleus and HSP90 marks the cytoplasm. F) Quantification of the band intensities seen in (E). G) Heatmap showing the expression levels of markers of the six different layers of the human cortex in week 15 organoids from deepest to the most superficial: TBR1, BCL11B (CTIP2), SATB2, POU3F2 (BRN2), CUX1, RELN. H) qPCR for markers of the six different layers of the human cortex in week 15 organoids *(WT: n=4, KO: n=4).* Data are represented as mean ± SEM. I) IF staining in week 15 COs validating the decreased levels of the deep-layer cortical marker CTIP2 (BCL11B) and superficial-layer marker SATB2 *(WT W15: n=3, KO 15: n=4)*.

Gene-set Enrichment Analysis (GSEA) and Gene ontology (GO) Enrichment analysis of the top 3,000 differentially expressed genes revealed, among others, inhibition of processes related to ATP synthesis coupled electron transport and oxidative phosphorylation (Figure 4B and Supplementary figure S4C), all of which are consistent with previous reported functions of WWOX in mouse models (Abu-Remaileh and Aqeilan, 2014, 2015; Abu-Remaileh *et al*., 2018). In addition, downregulation of genes related to negative regulation of cell cycle were seen, consistent with the previously reported diminished checkpoint inhibition (M. Abu-Odeh, Salah, *et al*., 2014; Abu-odeh, Hereema and Aqeilan, 2016). On the other hand, marked enrichment was seen in pathways related to regionalization, neuron fate commitment and specification, axis specification (Ventral-Dorsal & Anterior-Posterior) and glycolysis and gluconeogenesis, some of which are also supported by past studies (Wang *et al*., 2012; Abu-Remaileh and Aqeilan, 2014). As could be anticipated, upregulated genes were related to the development pathways such as Wnt pathway *(e.g. WNT1, WNT2B, WNT3, WNT3A, WNT5A, WNT8B, LEF1, AXIN2, GBX2, ROR2, LRP4, NKD1, IRX3, CDH1*) and the Shh pathway (e.g *SHH, GLI1, LRP2, PTCH1, HHIP, PAX1, PAX2)*.

Since WWOX has been previously implicated in the Wnt signaling pathway (Bouteille *et al*., 2009; Wang *et al*., 2012; Abu-Odeh, Bar-Mag, *et al*., 2014; Cheng *et al*., 2020; Khawaled *et al*., 2020), we set to further explore this in our COs models. First, we used our RNA-seq data to check the expression levels of different parts of the pathway - WNT family members (such as *WNT1, WNT3, WNT5A, WNT8B*), canonical targets (such as *Axin2, TCF7L2, LEF1, TCF7L1*), brain-specific targets (*IRX3, ITGA9, GATA2, FRAS1, SP5*) and receptors (*ROR2, FZD2, FZD10, FZD1*) (Figure 4C). We next validated some of these genes using qPCR (Figure 4D). Additionally, to prove Wnt activation, we demonstrated β-catenin translocation into the nucleus - a hallmark of the canonical Wnt-pathway. To this end, week 16 COs were sub-fractionated into cytoplasmic and nuclear fractions and immunoblotted (Figures 4E and 4F). We found a 1.2-1.5-fold increase in the normalized intensity of β-catenin in the nucleus of WWOX-KO COs, supporting the notion of Wnt-pathway activation after loss of WWOX. This was also supported by chronic activation of Wnt genes and targets in KO COs contrast to what is observed in WT COs (Supplementary figure S4E). Furthermore, examination of RNA expression levels of Wnt-related genes in W-AAV COs revealed downregulation of some genes, including *WNT3, WNT3A, WNT8B, WNT1, ROR2* (Supplementary figure S4H).

Recent evidence has demonstrated that activation of Wnt in forebrain organoids during development causes a disruption of neuronal specification and cortical layers formation (Qian *et al*., 2020). To address whether this occurs in WWOX-KO COs, we examined the expression levels of cortical layers markers (Qian *et al*., 2016) in our RNA-seq data (Figure 4G). Interestingly, changes were observed in all six-layers, with layers I-IV (marked by *TBR1, BCL11B, SATB2, POU3F2*) showing decreased expression, while superficial layers V-VI (marked by CUX1 and RELN) exhibiting marked increase. This pattern was also confirmed by qPCR (Figure 4H). Intriguingly, when we examined protein levels using immunofluorescent staining, we observed also impaired expression patterns and layering, with CTIP2^+^ (BCL11B) and SATB2^+^ neurons intermixing in WWOX-KO COs (Figure 4I). This defect was progressive, worsening at week 24 (Supplementary figures 4E). Surprisingly, when examining the effect of ectopic WWOX expression, a less clear phenotype was observed; although CTIP2^+^ cells and SATB2^+^ cells numbers recovered and layering improved in W-AAV COs compared to WWOX-KO COs (Supplementary figure S4H), RNA levels did not (Supplementary figure S4J). In contrast, expression of the superficial layers markers CUX1 and RELN (REELIN), that was upregulated in WWOX-KO, decreased in W-AAV, together with the upper layer marker POU3F2 (BRN2).

Overall, RNA-seq reveled impaired spatial patterning, axis formation and cortical layering in WWOX-KO COs, which is correlated with disruption of cellular pathways and activation of Wnt signaling. The reintroduction of WWOX prevents these changes to some extent, further supporting its possible implication in gene therapy.

### WWOX-Related Epileptic Encephalopathy (EIEE28) forebrain organoids presented similar phenotypes to WWOX-depleted COs

Although disease modeling using CRISPR-edited cell is a widely used tool, critiques argue against it for not modeling the full genetic background of the human patients. Therefore, we reprogramed peripheral blood mononuclear cells (PBMCs) donated from two families with WWOX-related diseases, differing in its severity: The first family, carries a c.517-2A>G splice site mutation (Weisz-Hubshman *et al*., 2019), resulting in the WOREE syndrome (EIEE28) phenotype in the homozygous patient (referred as WSM family) (Supplementary figures S5A and S5B). The second family carries a c.1114G>C (G372R) mutation (Mallaret *et al*., 2014) that results in the SCAR12 phenotype in the homozygous patients (referred as WPM family) (Supplementary figures S6A and S6B). All the iPSCs lines showed normal morphology for primed hPSCs, self-renewal capabilities (data not shown) and were evaluated for expression of pluripotent markers (Supplementary figures S5C, S5D, S6C and S6D). Since a major part of the phenotype was observed in the cortical part of the COs, we decided to employ a cortex-specific protocol, and generate forebrain organoids (FOs) (Qian *et al*., 2016, 2018).

First, we generated FOs from the healthy father and sick son of the WSM family (WSM F1 and WSM S5, respectively). FOs of both father and son exhibited similar morphology and growth throughout the protocol. Similar to what has been seen in the COs, staining of week 6 FOs showed WWOX expression localized to the VZ of WSM F1, though the expression was remarkably low and hard to visualize (Figure 5A) (Supplementary figure S6A). Although TUJ1^+^ positive cells and SOX2^+^ positive cells were comparable in numbers, WSM S5 showed no detectable levels of WWOX.

**Figure 5.**
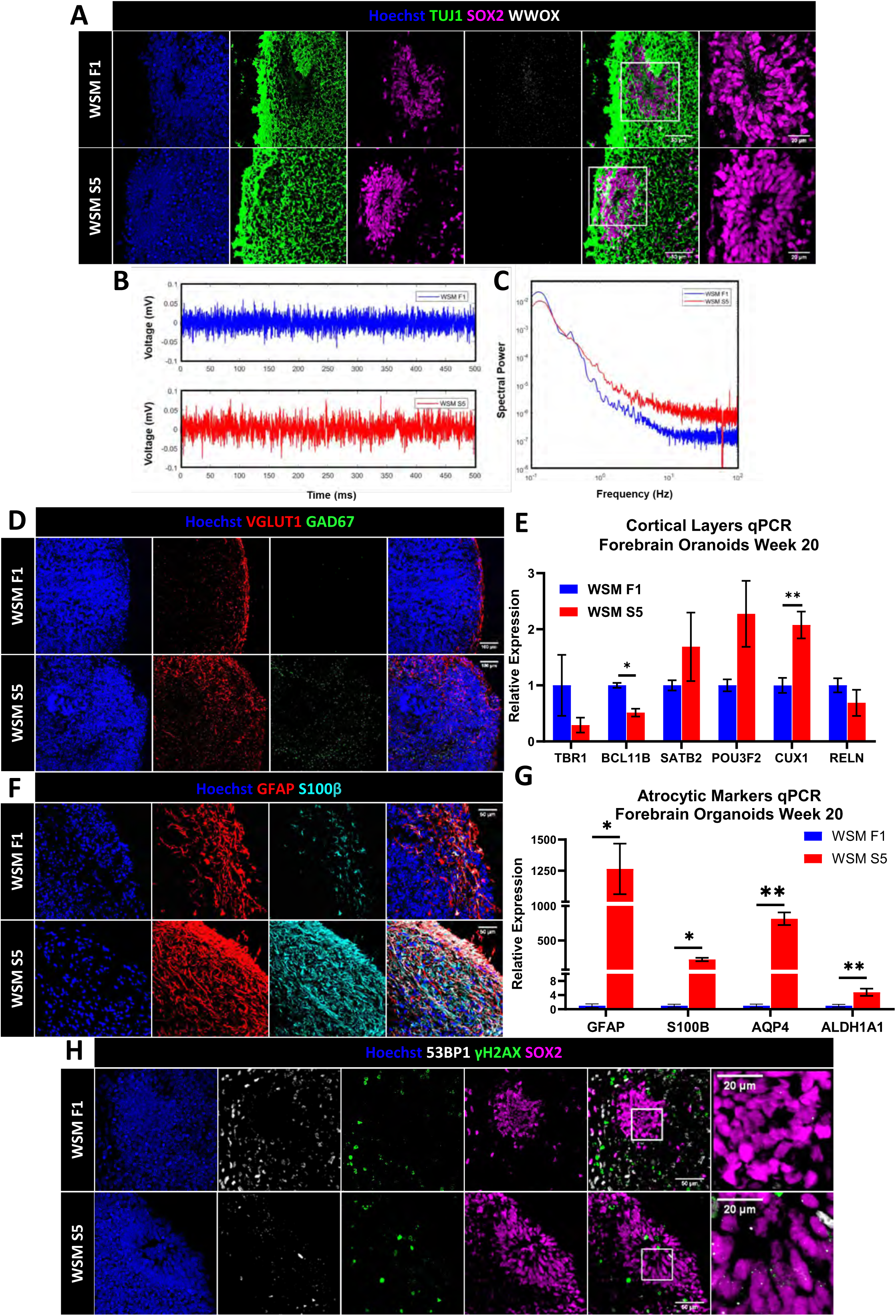
WWOX-Related Epileptic Encephalopathy Forebrain Organoids Presented Similar Phenotype to WWOX-KO COs. Peripheral blood mononuclear cells (PBMCs) were isolated from a WOREE patient and from his healthy parents and were reprogrammed into iPSCs, and subsequently were differentiated into forebrain organoids. A) Week 6 FOs of the healthy, heterozygote father (WSM F1) and his sick homozygote son (WSM S5) stained for WWOX expression. Similarly to WT COs, WWOX is expressed by the SOX2+ cells of the VZ in WSM F1 FOs, although barely visualized, with no detectable expression of WWOX in FOs from WSM S5. B) Sample recordings of week-12 FOs. C) Resulting spectral power graph from 12-week old FOs sample recordings. D) Immunofluorescent staining for VGLUT1 and GAD67 (GAD1) *(WSM F1: n=4, WSM S5: n=2)*. E) qPCR for the measurement of expression levels of cortical markers in 20 weeks FOs *(WSM F1: n=4, WSM S5: n=3).* Data are represented as mean ± SEM. F) Week 20 FOs stained for the astrocytic markers GFAP and S100β in week 20 WSM S5 FOs compared to the age matched WSM F1 FOs (WSM F1: *n=4, KO: n=2)*. G) qPCR quantifying the transcript levels of astrocytic markers in week 20 FOs *(WSM F1: n=4, WSM S5: n=3).* Data are represented as mean ± SEM. H) Week 6 FOs stained for the DNA damage markers γH2AX and 53BP1 under physiological conditions *(WSM F1: n=2, WSM S5: n=2).* For quantification, see Supplementary figure S5F.

Next, to evaluate whether WSM S5 FOs showed hyper-excitability, we performed LFP recordings of FO slices at week 12. Sample traces show an increase in amplitude of the WSM S5 slices compared to WSM F1 (Figure 5B). The power spectral density of traces further demonstrates this increase in amplitude in all frequencies above 0.5 Hz (Figure 6C).

Consistent with our findings in WWOX-KO COs, week 20 FOs showed elevated levels of GAD67 in WSM S5 compared to the healthy WSM F1 and similar expression of VGLUT1 (Figure 5D). Evaluation of expression levels of cortical layers’ markers showed a clear disruption in week 20 WSM S5 FOs compared to WSM F1 FOs, with diminished expression of deep-layer markers (TBR1 and BCL11B) and increased superficial-layers markers (SATB2, POU3F2 and CUX1). We further studied this phenotype by assessing expression levels of Wnt-related genes and found an increase in several Wnt-family members (*WNT1, WNT2B, WNT5A, WNT8B*) and Wnt-target genes (*Axin2, TCF7, TCF3*) (Supplementary figure S5E), suggesting Wnt-pathway activation.

Furthermore, similarly to COs, increased astrocytic marker levels were seen in week 20 WSM S5 FOs, both by immunostaining and qPCR (Figures 5F and 5G). Additionally, we observed elevated numbers of γH2AX and 53BP1 foci in the VZs of WSM S5 FOs compared to the healthy control (Figures 5H, quantified in Supplementary figure S6F), with a mean of 1.46 γH2AX foci/nuclei [95% CI=0.72-2.2] and 0.81 53BP1 foci/nuclei [95% CI=0.37-1.26] in WSM S5, compared to 0.63 γH2AX foci/nuclei [95% CI=0.45-0.8] and 0.3 53BP1 foci/nuclei [95% CI=0.17-0.43] in WSM F1. Together, our findings imply that WWOX-KOs successfully modeled the disturbances in development seen in FOs from WOREE patient further strengthening the model’s ability to recapitulate the patient’s disease.

Our next step was to study whether the WPM family, whose patients have a milder phenotype, present with similar phenotypes to WWOX-KO COs and WSM FOs. We generated FOs from the healthy heterozygous father and mother (WPM F2 and WPM M3) and their affected homozygous daughter and son (WPM D1 and WPM S1). As expected, FOs were indistinguishable in term of morphology, growth and expression of TUJ1 and SOX2 (Supplementary figure S6E), but while in the VZ of WPM F2 and WPM M3 WWOX was strongly detected, barely any signal was observed in WPM D1 and S1, consistent with WWOX levels in the iPSCs (Supplementary figure S6A). Surprisingly, transcript expression levels of neuronal markers did not show any clear difference in the ratio between glutamatergic and GABAergic neurons (Supplementary figure S6F). Although some differences were seen between FOs from lines with similar genotypes, the comparable levels of cortical layers’ marker expression between the healthy iPSCs lines (WPM F2 and WPM M3) and the disease bearing lines (WPM D1 and WPM S1) supported the notion of normal neuronal and cortical development (Supplementary figure S6G). Conversely, RNA levels of Wnt-genes did show a pattern suggestive of the Wnt-pathway activation (Supplementary figure S6H), which raises a question regarding its role in the pathogenesis of the milder disease. Immunostaining for astrocytic levels did not reveal any significant difference as well (Supplementary figure S6I), which was supported by RNA levels measurement (Supplementary figure S6J). Lastly, upon analyzing the DDR signaling in the FOs’ VZ, we did not observe significant differences in accumulation of DNA damage foci between healthy and sick SCAR12 individuals (Supplementary figure S6K).

## Discussion

EIEEs are a group of severe neurologic syndromes whose underlying molecular pathology is unknown. Together with the lack of accessibility of human samples, it not surprising that the current medical treatment is lacking. Our study set out to utilize the major technological advances in developmental biology, together with the role of WWOX in the severe WOREE syndrome, to model human refractory EIEEs in a tissue-relevant context. By utilizing genetic and epigenetic editing tools along with electrophysiology, we observed hyper-excitability in both WWOX CRISPR-edited COs and patient-derived FOs, therefore successfully demonstrating epileptiform activity. Organoid slices were particularly active in the lower frequency ranges – an attribute generally characteristic of seizure-like activity (Haddad *et al*., 2014). We then further examined the cellular and molecular changes highlighting possible mechanisms for the disease pathophysiology. First, although the neuronal population was largely intact in terms of quantity, we noticed a marked increase in GABAergic markers. This finding is even more surprising when considering the decrease in GABA-receptor components seen by RNA-seq. This can indicate a disruption in development of normal and balanced neuronal networks, supporting the increased electrical activity observed in these organoids. It should be noted that several lines of evidence implicate that during development, GABAergic synapses have a depolarizing effect (Obata, Oide and Tanaka, 1978; Ben-Ari *et al*., 2007; Murata and Colonnese, 2020). Seizure dynamics in developmental epilepsies are known to be dependent on depolarizing GABA responses, particularly due to an accumulation of intracellular chloride resulting in a depolarized chloride reversal potential, thereby causing increased excitability, instead of hyperpolarization upon activation of GABA_A_ receptors (Khalilov *et al*., 2005; Ben-Ari *et al*., 2007). The evidence of increased mean spectral power in WWOX-depleted COs and WSM FOs, and its recovery in the presence of lentivirus containing WWOX, further strengthens the idea that depolarizing GABA plays a key role in seizure susceptibility. These findings shed a new light on the lack of efficacy of common anticonvulsant therapies on immature neurons (Khalilov *et al*., 2005; Murata and Colonnese, 2020) – making WWOX-depleted COs a useful model to test and study novel therapies targeting excitatory GABAergic responses.

Secondly, we closely examined other populations seen in brain organoids, and found an increase in astrocytic markers, while the RGs population, which express high levels of WWOX, seemed to maintain normal proportions. This pattern was detected early on and appeared to stem from the RGs of the VZ themselves, and not from the APCs. A possible explanation is the impaired DDR signaling observed in WWOX-depleted organoids; Previous studies in both ESCs-derived and primary murine neural stem cells (NSCs) found that accumulation of DNA damage foci, either in the nuclear or mitochondrial DNA, causes NSCs to astrocytic differentiation (Wang *et al*., 2011; Schneider *et al*., 2013). In the CNS, physiological DNA breaks, can form by replicative stress (mainly in dividing progenitor cells), by oxidative and metabolic stress as a result of accumulation of reactive oxygen species (ROS) and even by neuronal activity (as part of developmental processes and learning) (Suberbielle *et al*., 2013; Madabhushi, Pan and Tsai, 2014; Madabhushi *et al*., 2015). Impaired repair of these breaks is linked with CNS pathology and neurodegeneration (Suberbielle *et al*., 2013; Madabhushi, Pan and Tsai, 2014; Shanbhag *et al*., 2019). Our findings suggest a homeostatic role for WWOX in the RGs of the VZ, in which WWOX maintains proper DDR signaling in physiological conditions and prevents accumulation of DNA damage associated with impaired differentiation.

Although the ability of brain organoids to develop functional synapses and complex neural network dynamics is rapidly being established through intensive research (Trujillo *et al*., 2019; Sidhaye and Knoblich, 2020), the capability to model epileptiform activity is only recently being studied (Samarasinghe *et al*., 2019; Sun *et al*., 2019). Sun et al., (2019) utilized brain organoids to model Angelman syndrome using UBE3A-KO hESCs, recapitulating hyperactive neuronal firing, aberrant network synchronization and the underlying channelopathy which was observed in 2D and mouse models. Samarasinghe et al. (2019) took advantage of the organoid fusion method and generated organoids enriched with inhibitory interneurons from a Rett syndrome patient’s iPSCs. In the disease-bearing organoids, they observed susceptibility for hyperexcitability, reductions in the microcircuit clusters, recurring epileptiform spikes and altered frequency oscillations, which was traced back to dysfunctional inhibitory neurons. Furthermore, the model was used to test treatment options by treating the mutated organoids with Valproic acid (VPA) or with the TP53 inhibitor, Pifithrin-α (PFT), showing improved neuronal activity compared to the treatment with vehicle, with better results using PFT rather than VPA. Although pioneering, these studies focused on the electrophysiological changes seen in the disease-modeling organoids. Considering the lack of gross neurohistological changes in epileptic patients to direct the mechanistic research (Blumcke *et al*., 2017), our study sought to strengthen the utilization of brain organoids for the molecular study of epilepsy. This end was highlighted by bulk RNA-seq analysis, showing defective regional identity acquisition, cortical layer disruption and Wnt signaling activation. The latter is of particular interest in light of the purposed role for Wnt signaling pathway as a regulator of seizure-induced brain consequences, and therefore a possible target for treatment (Yang *et al*., 2016; Qu *et al*., 2017; Hodges and Lugo, 2018).

In agreement with our findings, a recent study that examined the brain histology of a fetus suffering from the WOREE syndrome, reported anomalous migration of the external granular layer within the molecular layer of the cortex, a phenotype that was validated also in a rat model with spontaneous WWOX mutations (Iacomino *et al*., 2020). A recent study of *Wwox-*null mice demonstrated that activation of the Wnt/β-catenin signaling through use of GSK3β inhibitors suppressed PTZ-induced epileptic seizures, highlighting it’s possible role in its pathogenesis (Cheng *et al*., 2020). Other known binding partners are the Disheveled proteins Dvl1/2, with the latter being inhibited by WWOX, therefore attenuating the Wnt-pathway (Bouteille *et al*., 2009; Abu-Odeh, Bar-Mag, *et al*., 2014). Our study further highlights a possible cross talk between Wnt-activation and DNA damage, a phenomenon that was previously described (Elyada *et al*., 2011). This is very much in-line with the previously described pleiotropic functions of WWOX (Abu-Remaileh *et al*., 2015) and with the reduced negative regulation of cell cycle and MDM2 levels seen in our RNA-seq. We found accumulation of DNA breaks in Ki67^+^ cells in the VZ of KO COs, which might be explained by Wnt activation, promoting proliferation and likely replicative stress.

In addition to disease modeling in brain organoids, we attempted to rescue the phenotypes seen by re-introducing WWOX to the hESCs genome. This resulted in supraphysiological expression of WWOX in all cell populations seen in COs and a partial rescue. These results provide a proof-of-concept for successful reintroduction of WWOX as a mean of therapeutic intervention. Yet, our findings suggest the importance of optimizing population-targeted delivery and fine tuning of expression levels for successful genetic therapy approaches in WOREE patients.

Lastly, we generated FOs from patients suffering from the relatively milder phenotype - SCAR12. Our results indicate that SCAR12 FOs do not suffer from the same developmental abnormalities as the WOREE patient. SCAR12 FOs exhibited very mild, if any, differences in the forebrain neuronal population development, astrocytes development and DDR signaling. This strengthens the system’s ability to model the differences seen between the syndromes, and points out the need of closer examination of the rare SCAR12 syndrome and the pleiotropic functions of WWOX (Abu-Remaileh *et al*., 2015). It is noteworthy that although there is a marked difference in WWOX-expression in the healthy heterozygote parents from different families, there is a very minor difference in levels observed in the affected homozygote patients (Figure 6A and supplementary figures S5A, S6A and S6E). These results raise the question whether the disease severity is correlated with the functional levels of WWOX rather than its the total expression levels.

Overall, our data demonstrate the ability of brain organoids to model childhood epileptic encephalopathies, while elucidating the pathological changes seen in patients with germline mutations of WWOX and possible approaches for treatment development.

## Limitations of Study

As samples from EIEE-patients in general, and WWOX-related disorders in particular, are limited, we generated patient-derived organoids from only two families, one of each syndrome. This makes generalizing our results more difficult, a problem we partially addressed by gene-manipulation in hESCs. Another limitation stems from the well-described heterogeneity of brain organoids which we dealt with by analyzing several repeats and confirmed in patient-derived models.

## Acknowledgments

We would like to thank all members of the Aqeilan’s lab for fruitful discussion. We are grateful to Dr. Abed Nasereddin and Dr. Idit Shiff from the Genomic Core Facility for their help. The Aqeilan’s lab is funded by the European Research Council (ERC) [No. 682118], Proof-of-concept ERC grant [No. 957543] and the KAMIN grant from the Israel Innovation Authority [No. 69118].

## Author Contributions

Conceptualization, D.J.S., S.R. and R.I.A.; Methodology, D.J.S. S.R., J.H.H., and R.I.A.; Investigation, D.J.S. and A.S.; Writing – Review & Editing, D.J.S., A.S., P.L.C., and R.I.A.; Funding Acquisition, R.I.A.; Resources, E.M., M.M. and J.H.H; Project Administration, R.I.A.; Supervision, P.L.C. and R.I.A.

## Declaration of interests

The authors declare no competing interests.

## Materials and Methods

### Cell culture and plasmids

WiBR3 hES cell line and the generated iPS cell lines were maintained in 5% CO_2_ conditions on irradiated DR4 mouse embryonic fibroblasts (MEF) feeder layers in FGF/KOSR conditions: DMEM-F12 (Gibco; 21331-020 or Biological Industries; 01-170-1A) supplemented with 15% Knockout Serum Replacement (KOSR, Gibco; 10828-028), 1% GlutaMax (Gibco; 35050-038), 1% MEM non-essential amino acids (NEAA, Biological Industries; 01-340-1B), 1% Sodium-pyruvate (Biological Industries; 03-042-1B), 1% Penicillin-Streptomycin (Biological Industries; 03-031-113), and 8ng/mL bFGF (Peprotech; 100-18B). Medium was changed daily and cultures were passaged every 5– 7 days either manually or by trypsinization with Trypsin type C (Biological Industries; 03-053-1B), and Rho-associated kinase inhibitor (ROCKi, also known as Y27632) (Cayman; 10005583) was added for the first 24-48h at a 10µM concentration.

For transfection of hESCs, cells were cultured in 10µM ROCKi 24h before electroporation. Cells were detached using Trypsin C solution and resuspended in PBS (with Ca^2+^ and Mg^2+^) mixed with a total of 100μg DNA constructs, and electroporated in Gene Pulser Xcell System (Bio-Rad; 250 V, 500μF, 0.4cm cuvettes). Cells were subsequently plated on MEF feeder layers in FGF/KOSR medium supplemented with ROCKi. For WWOX-KO, px330 plasmid containing the sgRNA targeting exon 1 was co-electroporated in 1:5 ratio with pNTK-GFP, and 48hr-later, GFP-positive cells were sorted and subsequently plated sparsely (2,000 cells per 10cm plate) on MEF feeder plates for colonies isolation, ∼10 days later. For WWOX-reintroduction, pAAVS-2aNeo-UBp-IRES-GFP plasmid cloned to carry the WWOX coding sequence was co-electroporated with px330 targeting the AAVS locus (Guernet *et al*., 2016), sorted for GFP and selected with 0.5µg/ml Puromycin for colonies isolation. Gene-editing was validated via Western Blot. sgRNA sequences are noted in supplementary table 3.

For RNA or protein isolation, hPSCs were passaged onto Matrigel-coated plates (Corning; 356231) as indicated above and were cultured in NutriStem hPSC XF medium (Biological Industries; 05-100-1A).

### Cerebral organoid generation, culture, and lentiviral infection

Cerebral organoids were generated from hESCs as previously described(Lancaster *et al*., 2013, 2018; Lancaster and Knoblich, 2014; Bagley *et al*., 2017), with the following changes:

Human WiBR3 cells were maintained on mitotically inactivated MEFs. 4-7 days before protocol initiation, cells were passaged onto 60mm plates coated with either MEFs or Matrigel (Corning; FAL356231) and grown until 70-80% confluency was reached. On day 0, hESCs colonies were detached from MEFs with 0.7mg/ml collagenase D solution (Sigma; 11088858001) and dissociated to single cell suspension using a quick two minutes treatment with Trypsin type C. For cells cultured on Matrigel, collagenase D treatment was skipped, and cells were immediately dissociated with trypsin type C, with no other variations in protocols from this point forward. Although only empirically observed, no major differences were seen in final outcome, however MEF-cultured hESCs seemed to have better success rates of neural induction and therefore were preferentially used.

After dissociation, cells were counted and suspended in hESCs medium, composed of DMEM/F12 supplemented 20% KOSR, 3% USDA certified hESCs-quality FBS (Biological Industries), 1% GlutaMax, 1% NEAA, 100µM 2-mercaptoethanol (Sigma; M3148), 4ng/ml bFGF 6and 10µM Rocki, sterilized through 0.22μm filter, and 9,000 cells were seeded in each well of an ultra-low attachment 96 v-well plates (S-Bio Prime; MS-9096VZ) for embryoid bodies (EBs) formation. EBs were fed every other day for another 5 days, in which fresh bFGF and ROCKi were added in the first change. At day 6, the medium was replaced with Neural Induction (NI) medium(Bagley *et al*., 2017), composed of DMEM/F12, 1% N2 supplement (Gibco; 17502048), 1% GlutaMax, 1% MEM-NEAA, 1µg/ml Heparin solution (Sigma; H3149) sterilized through 0.22μm filter. NI medium was changed every other day until establishment of neuroepithelium (usually on days 11-12), were quality control was performed as indicated(Lancaster and Knoblich, 2014; Bagley *et al*., 2017), and well-developed EBs were embedded in Matrigel droplets(Lancaster and Knoblich, 2014; Bagley *et al*., 2017). Droplets were transferred to 90mm sterile, non-treated, culture dishes (Miniplast; 825-090-15-017) with Cerebral Differentiation Medium (CDM) composed of 1:1 mixture of DMEM/F12 and Neurobasal medium (Gibco; 21103049 or Biological Industries; 06-1055-110-1A), 0.5% N2 supplement, 1% B27 supplement without vitamin A (Gibco; 12587010), 1% GlutaMax, 1% penicillin/streptomycin, 0.5% NEAA, 50µM 2-mercaptoethanol, 2.5µg/ml human recombinant Insulin (Biological Industries; 41-975-100) and 3µM CHIR-99021 (Axon Medchem ; 1386) sterilized through 0.22μm filter. Medium was changed every other day. From day 16 onward, organoids were cultured on an orbital shaker at 37°C and 5% CO_2_ in Cerebral Maturation Medium (CMM)(Lancaster *et al*., 2018) composed similarly to CDM, with B27 supplement changed to B27 supplement containing vitamin A (Gibco; 17504044), without CHIR-99021, and containing 400ul mM vitamin C (Sigma; A4403) and 12.5mM HEPES buffer (Biological Industries; 03-025-1B). Medium was changed every 2-4 days. From week 6, 1% Matrigel was added to the medium. To reduce chances of contamination, every 30 days the organoids were moved to fresh sterile plates. For all analysis, organoids from the same batch were used, unless stated otherwise.

Lentiviral transduction of WWOX was carried as previously published(Deverman *et al*., 2016; Khawaled *et al*., 2019). Briefly, Viruses carrying WWOX were generated from pDEST12.2TM destination vector (Gateway Cloning Technology). After ultracentrifugation, titer was determined empirically by infecting 293T cells. At day 35 of culture, individual COs were transferred to an Eppendorf tube containing CMM medium with 1:100 of virus containing medium and 5µg/ml Polybrene (Merck; TR-1003-6) and incubated over-night. The day after, organoids were put back on shaking culture with fresh medium.

### Reprogramming of somatic cells

Blood samples from WOREE and SCAR12 families were donated under the approval of the Kaplan Medical Center Helsinki committee for research purposes only. Derivation of iPSCs directly from PBMCs was conducted by infection with the Yamanaka factors and Sendai virus Cyto-Tune-iPS2.0 Kit according to manufacturer’s instructions.

Briefly, blood samples. PBMCs were isolated by ficoll gradient and were cultured with StemPro-34™ medium (Gibco; 10639-011) supplemented with StemPro-34 Nutrient Supplement (Gibco; 10639-011), 100ng/ml human SCF (Peprotech; 300-07), 100ng/ml human FLT-3 ligand (R&D Systems; 308-FKE), 20ng/ml human IL-3 (Peprotech; 200-03) and 10ng/ml Human IL-6 (Peprotech; 200-06). After 24hr, half of the medium was replaced. After additional 24hr, day 0 of the protocol, cells were transferred for 6-well plates, reprogramming virus mixture was added, and the plate was centrifuged at 1000xg for 30 minutes at room temperature. Cells were re-suspended and placed back in the incubator overnight. The next day, to get rid of the remaining virus, the cells were centrifuged washed and re-suspended in fully supplemented StemPro-34 medium, with extra medium addition on day 2. On day 3, cells were transferred to 10cm MEF-coated plates, with half the medium replaced with complete StemPro-34 without cytokines and half medium changes every other day. By day 7 cells in different phases of reprogramming were seen, and the medium was gradually changed into mTeSR supplemented with 10µM ROCKi to prevent reprogramming-related apoptosis. On day 16, colonies with normal morphology and growth rate were picked, expanded, validated for expression of pluripotency markers, and sequenced for WWOX-mutations.

### Forebrain organoid generation and culture

Forebrain organoids were generated from iPSCs as previously described(Qian *et al*., 2016, 2018), with the changes noted below: iPSCs cells were maintained on mitotically inactivated MEFs. 4-7 days before protocol initiation, cells were passaged onto MEF-coated 6-well mm plates and were cultured up to 70-80% confluency. On day 0, iPSCs colonies were detached, dissociated, and counted same as for COs, and resuspended in hPSCs medium containing DMEM/F12, 20% KOSR, 1% GlutaMax, 1% MEM-NEAA, 1% penicillin/streptomycin and 100µM 2-mercaptoethanol, and 9,000 cells were seeded in 96 v-well plate. On day 1, medium was changed to Neuroectoderm Medium (NEM) which is hPSCs medium freshly supplemented with 2µM A83 (Axon Medchem; 1421) and 100nM LDN-193189 (Axon Medchem; 1527), which was changed every other day. On days 5 and 6, half of the medium was aspirated and replaced by Neural Induction Medium (NIM) composed of DMEM/F12, 1% N2 supplement, 1% GultaMax, 1% Penicillin/Streptomycin, 1% NEAA, 10µg/ml Heparin, 1µM CHIR-99021 (Axon Medchem ; 1386) and 1µM SB-431542 (Sigma; S4317). On day 7, quality control and Matrigel embedding was performed as indicated(Qian *et al*., 2018), and EBs were continued to be cultured in NIM with medium changes every other day. At day 14 Matrigel removal was preformed(Qian *et al*., 2018), medium was changed to Forebrain Differentiation Medium (FDM) composed of DMEM/F12, 1% N2 Supplement, 1% B27 with vitamin A, 1% NEAA, 1% GlutaMax, 1% Penicillin/Streptomycin, 50µM 2-mercaptoethanol and 2.5µg/ml Insulin, and transferred to an orbital shaker at 37°C and 5% CO_2_. Medium was changed every 2-3 days. On day 71, medium was changed to Forebrain Maturation Medium (FMM), containing Neurobasal medium, 1% B27 supplement with vitamin A, 1% GlutaMax, 1% Penicillin/Streptomycin, 50µM 2-mercaptoethanol, 200µM Vitamin C, 20 ng/ml human recombinant BDNF (Peprotech; 450-02), 20ng/ml human recombinant GDNF (Peprotech; 450-10), 1µM Dibutyryl cAMP (Sigma; D0627), 1ng/mL TGF-β1 (Peprotech; 100-21C). Medium was changed every 2-3 days.

### Immunofluorescence

Organoids fixation and immunostaining were performed as previously described(Mansour *et al*., 2018). Briefly, organoids were washed three times in PBS, then transferred for fixation in 4% ice-cold paraformaldehyde for 45 min, washed three times in cold PBS, and cryoprotected by over-night equilibration in 30% sucrose solution. The next day, organoids were embedded in OCT, snap frozen on dry ice and sectioned at 10μm by Leica CM1950 cryostats.

For immunofluorescent staining, sections were warmed to room temperature and washed in PBS for rehydration, permeabilized in 0.1% Triton X in PBS (PBT), and then blocked for 1hr in blocking buffer containing 5% normal goat serum (NGS), 0.5% BSA in PBT. The sections were then incubated at 4°C overnight with primary antibodies diluted in blocking solution. The day after, sections were then washed in 3 times while shaking in PBS containing 0.05% Tween-20 (PBST) and incubated with secondary antibodies and Hoechst33258 solution diluted in blocking buffer for 1.5hr. Slides were washed four times in PBST while shaking, and coverslips were mounted using Immunofluorescence Mounting Medium (Dako; s3023). Sections were imaged with Olympus FLUOVIEW FV1000 confocal laser scanning microscope and processed using the associated Olympus FLUOVIEW software. γH2AX-positive nuclei were manually counted using NIH ImageJ and statistically analyzed as later described.

List of primary and secondary antibodies used in this work, together with dilutions details can be found in Supplementary table 4.

### Electrophysiological Recordings

Organoids were embedded in 3% low temperature gelling agarose (at ∼36°C) and incubated on ice for 5 minutes, after which they were sliced to 400µm using a Leica 1200S Vibratome in sucrose solution (in mM: 87 NaCl, 25 NaHCO3, 2.5 KCl, 25 Glucose, 0.5 CaCl2, 7 MgCl2, 1.25 NaHPO4, 75 Sucrose) at 4°C. Slices were incubated in artificial cerebrospinal fluid (ACSF, in mM: 125 NaCl, 25 NaHCO3, 2.5 KCl, 10 Glucose, 2.5 CaCl2, 1.5 MgCl2; pH 7.38, 300mOsm) for 30 minutes at 37°C, followed by 1 hour at RT. During recordings, slices were incubated in the same ACSF at 37°C with perfused carbogen (95% O_2_, 5% CO_2_), in baseline condition. Local field potential (LFP) and whole-cell patch clamp recordings were done using electrodes pulled from borosilicate capillary glass and positioned 150µm deep from the outer rim of each slice (see Supplementary figure S2A). LFP electrodes were filled with ACSF, while patch electrodes were filled with internal solution. Data was recorded using MultiClamp software at a sampling rate of 25,000 Hz. Data was analyzed using MATLAB software. Traces were filtered using (1) 60 notch filter (with 5 harmonics) to eliminate noise and (2) 0.1 Hz high-pass IIR filter to eliminate fluctuations from the recording setup. The detrend feature (using the hamming window) was then used to eliminate large variations in the signal and the normalized spectral power was calculated using Fast-Fourier Transform. The area under the curve of the power spectral density plots was calculated by taking the sum of binned frequencies over specific frequency ranges.

### Immunoblot analysis and Subcellular Fractionation

For total protein, organoids homogenized in lysis buffer containing 50 mM Tris (pH 7.5),150 mM NaCl, 10% glycerol, and 0.5% Nonidet P-40 (NP-40) that was supplemented with protease and phosphatase inhibitors. For separation of cytoplasmic fraction, organoids were grinded in a hypotonic lysis buffer [10 mmol/liter HEPES (pH 7.9), 10 mmol/liter KCl, 0.1 mmol/liter EDTA] supplemented with 1 mmol/liter DTT and protease and phosphatase inhibitors. The cells were allowed to swell on ice for 15 min, and then 0.5% NP-40 was added, and cells were lysed by vortex. After centrifugation, the cytoplasmic fraction was collected. Afterwards, nuclear fraction was obtained by incubating remaining pellet in a hypertonic nuclear extraction buffer [20 mmol/liter HEPES (pH 7.9), 0.42 mol/liter KCl, 1 mmol/liter EDTA] supplemented with 1 mmol/liter DTT for 15 min at 4°C while shaking. The samples were centrifuged, and liquid phase was collected.

Western blotting was performed under standard conditions, with 40-50µg protein used for each sample. Blots were repeated and quantified 2–3 times per experiment in Bio-Rad’s Image Lab software. Representative images of those repeated experiments are shown.

### RNA extraction, reverse transcription-PCR, and qPCR

Total RNA was isolated using Bio-Tri reagent (Biolab; 9010233100) as described by the manufacturer for Phenol-Chloroform based method. 0.5-1µg of RNA was used to synthesize cDNA using a qScript cDNA Synthesis kit (QuantaBio; 95047). qRT-PCR was performed using Power SYBR Green PCR Master Mix (Applied Biosystems; AB4367659). All measurements were performed in triplicate and were standardized to the levels of either HPRT or UBC. All primer sequence used are noted in Supplementary table 5

### Library preparation and RNA-sequencing

Library preparation and RNA-sequencing was performed by the Genomic Applications Laboratory in the Hebrew University’s Core Research Facility following standard procedures. Briefly, RNA quality was assessed by using RNA ScreenTape kit (Agilent Technologies; 5067-5576), D1000 ScreenTape kit (Agilent Technologies; 5067-5582), Qubit(r) RNA HS Assay kit (Invitrogen; Q32852) and Qubit(r) DNA HS Assay kit (Invitrogen; 32854).

For mRNA library preparation, 1ug of RNA per sample was processed using KAPA Stranded mRNA-Seq Kit with mRNA Capture Beads (Kapa Biosystems; KK8421). Library was eluted in 20µl of elution buffer, adjusted to 10mM, then 10µl (50%) from each sample was collected and pooled in one tube. Multiplex samples Pool (1.5pM including PhiX 1.5%) was loaded in NextSeq 500/550 High Output v2 kit (75 cycles) cartridge (Illumina; FC-404-1005) and loaded on NextSeq 500 System machine (Illumina), with 75 cycles and single-read sequencing conditions.

For library quality control, Fastq files were tested with *FastQC* (ver.0.11.8) and trimmed for residual adapters, low-quality bases (Q=20) and read length (20 bases). Trimming was performed with *trim galore (*ver.0.6.1). Read counts were high around 30M-50M per sample and decreased negligibly after filtering. Transcriptome mapping was performed with *salmon* (ver.1.2.1) in its mapping-based mode, turning on both validate mapping mode and gc-bias correction. Prior to alignment, a salmon index was created based on HS GRCh38 CDNA release 99 (Nov 2019) using kmer size of 25. Salmon mapping reports both raw transcripts count and TPM counts. Resulting mapping rates is high between 80%-90%. A total of 8 COs samples were sequenced (4 WT COs, 4 KO COs) – one WT sample failed our preliminary quality control (low read count and low transcriptome mapping rate). Another WT sample did not cluster with any of the other samples (neither WWOX-KO nor WT) was apparent in both PCA and Dendrogram analysis (Data not shown). These two samples were extracted from further analysis, giving a total of 6 samples used for further analysis. For differentially expressed genes determination (KO vs WT), raw transcript counts were filtered for minimal overall count of 10 on all six samples and imported with R package *tximport* (ver.1.16.1) for analysis with *DEeq2* (ver.1.28.1). Counts were normalized by DESeq2 and differentially expressed genes were filtered, setting alpha to 0.01. Mean based fold change was calculated as well as a shrink-based fold change based on *apeglm* (ver.1.10.0). The resulting set of 15,370 genes is illustrated in a “volcano scatter plot” showing fold change against p-values (Supplementary figure S4B).

For heatmaps preparation shown in Figures 4A and 4C, the list of Differentially Expression Genes was separated to upregulated (WWOX-KO expression was higher than WT expression) and downregulated sub lists. Each sub-list was sorted by Fold Change values and top 100 genes were selected from each sub-list. For each of the selected genes, log2 normalized counts were scaled and presented in a heatmap form using heatmap.2 from R package *gplots* (ver.3.0.3).

For the heatmap seen in figure 4G, Log2 normalized counts for each of the six cortical layer gene markers were scaled and presented in a heatmap form using heatmap.2 from R package *gplots*. For Enrichment plot (Figures 4B and supplementary figure S4D), Gene set enrichment analysis was performed with Broad Institute *GSEA* software (ver.4.0.3). Input included 15,348 genes ranked by log2 of Fold Change. GO sets is Broad Institute set c5.all.v7.0. Permissible sets are those with at least 15 genes and no more than 500 genes. For Enrichment plot seen in supplementary figure S4C, Gene set enrichment analysis was performed with *WebGestalt*R (ver.0.4.3). Input includes 3000 Differentially Expressed genes with most significant adjusted p-values and ranked by log2 of fold change. Gene sets are GO Biological Processes. Permissible sets in this analysis are those with at least 10 genes and no more than 500 genes. PCA plot (Supplementary figure S4A) of first two components was calculated and plotted with base R functions. Calculation is based on log2 transformed and normalized counts adding pseudo count of 1.)

### Statistics

Results of the experiments were expressed as mean ± SEM. First, Wilks-Shapiro test was used to determine normality: For normally distributed samples, a two-tailed Student’s t-test with Welch’s correction was used to compare the values of the test and control samples. For non-normally distributed samples, the non-parametric Mann-Whitney test was used. For comparisons between more than two samples, one-way ANOVA was used, correcting for the multiple comparisons with Tukey’s multiple comparisons test. For the kinetics experiments (Supplementary figure S4E), the analysis was corrected for multiple t-tests using the Holm-Šídák method, without assuming equal SD. P-value cutoff for statistically significant results were used as following: *p ≤ 0.05, **p ≤ 0.01, ***p ≤ 0.001, ****p ≤ 0.0001. Statistical analysis and visual data presentation were preformed using GraphPad Prism 8.

## Data and Code Availability

The RNA-seq datasets generated during this study are available at GEO: GSE156243.

## Supplemental Figures’ Titles and Legends

**Figure S1.**
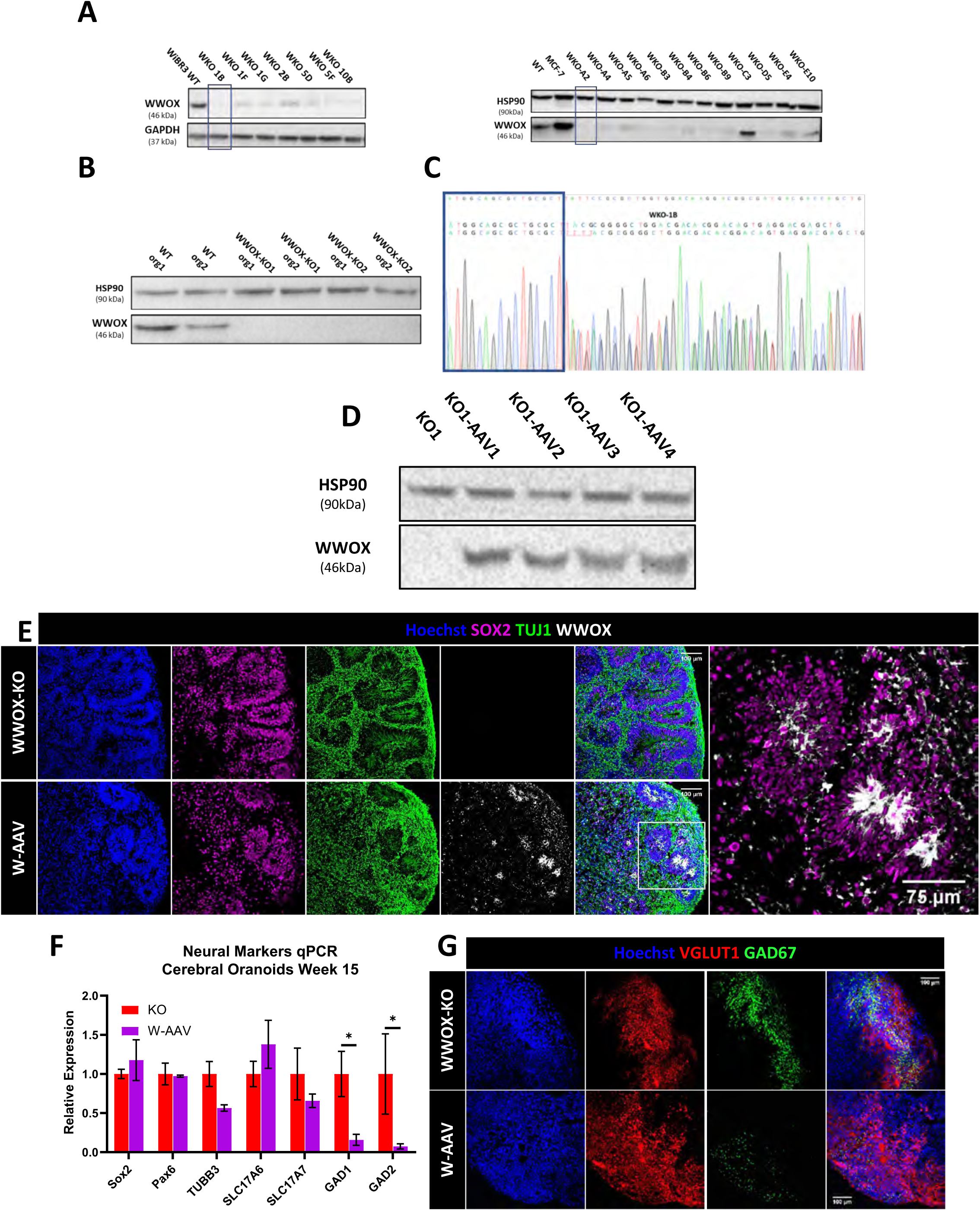
Generation and characterization of WWOX Knock-out Cerebral Organoids. A) Western blot (WB) analysis of WiBR3 hESCs individual colonies after CRISPR-editing targeted to exon 1 of the WWOX gene. Clone 1B (KO1) and clone A2 (KO2) were selected for the organoid generation. MCF-7 was used as a positive control highly expressing WWOX. B) Week 6 COs WB for WWOX-expression. C) Sanger sequencing of the WWOX KO clone 1B (WKO-1B). On one allele, an insertion of one nucleotide caused a frame shift, while in the other an insertion of 4 nucleotide occurred. Both resulted in a downstream premature stop codon (data not shown). D-G: WWOX-KO WiBR3 hESCs were introduced with a plasmid containing WWOX coding sequence and targeting the safe harbor locus AAVS. This results in hESCs over-expressing WWOX (WWOX-OE) under UBP promotor from the AAVS locus (W-AAV). D) Western blot analysis of KO1-hESCs clones introduced with WWOX coding sequence. E) Generation and validation of WWOX-expression in the W-AAV COs at week 10 (KO: n=3, W-AAV: n=4). F) qPCR assessment of expression levels of different neural markers in 15 weeks COs: Sox2 and Pax6 (progenitor cells), SLC17A6 and SLC17A (VGLUT2 and VGLUT1; glutamatergic neurons) and GAD1 and GAD2 (GAD67 and GAD65; GABAergic neurons). Y-axis indicated relative expression fold change. Data are represented as mean ± SEM (*KO: n=4, W-AAV: n=4)*. G) Staining for the glutamatergic neuron marker VGLUT1 and the GABAergic neuron marker GAD67 week 10 W-AAV COs and WWOX-KO COs (KO: n=3, W-AAV: n=4).

**Figure S2.**
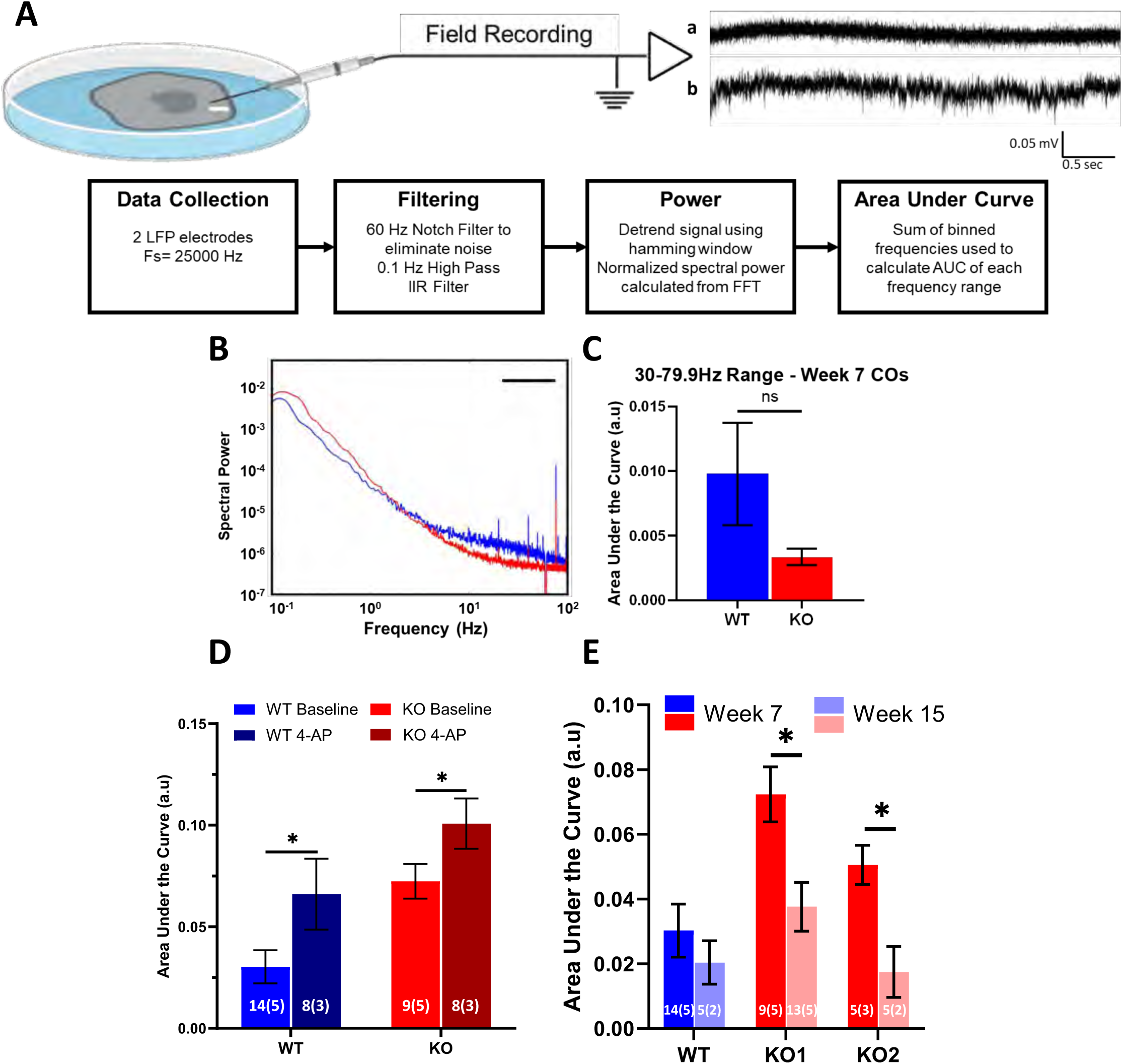
WWOX-KO Cerebral Organoids Demonstrated hyper-excitability and epileptiform activity. Sample recordings from 7-week old hESC-derived Cerebral Organoids (COs). (A) Top panel: Schematic of organoid slice set up. A borosilicate glass electrode is used for local field potential recordings (LFP) and is positioned 150µm from the edge of the slice (indicated by white bar). A sample recording is shown (a) before and (b) after administration of 100µM 4-AP. Bottom panel: Flowchart of steps used for signal processing field recordings. (IIR = Infinite Impulse Response, FFT = Fast Fourier Transform, AUC = Area Under Curve). (B) Mean spectral power of WT and 2 KO lines at week 7 in baseline conditions. (C) Normalized area under the curve of the mean spectral power in (A) for the low gamma range (30-79.9 Hz). Data represented by mean ± SEM. The two-tailed unpaired Student’s t-test was used to test statistical significance. The numerals in all bars indicate the number of analyzed slices and organoids (i.e. slices (organoids)). (D) Normalized area under the curve of the mean spectral power of WT and KO lines at week 7 in baseline and 100µM 4-AP conditions, for the 0.25-1 Hz frequency range. The two-tailed unpaired Student’s t-test was used to test statistical significance. The numerals in all bars indicate the number of analyzed slices and organoids. (E) Normalized area under the curve of the mean spectral power WT and 2 KO lines at week 7 and 15 in baseline conditions, for the 0.25-1 Hz frequency range. The two-tailed unpaired Student’s t-test was used to test statistical significance. The numerals in all bars indicate the number of analyzed slices and organoids (i.e. slices (organoids)).

**Figure S3.**
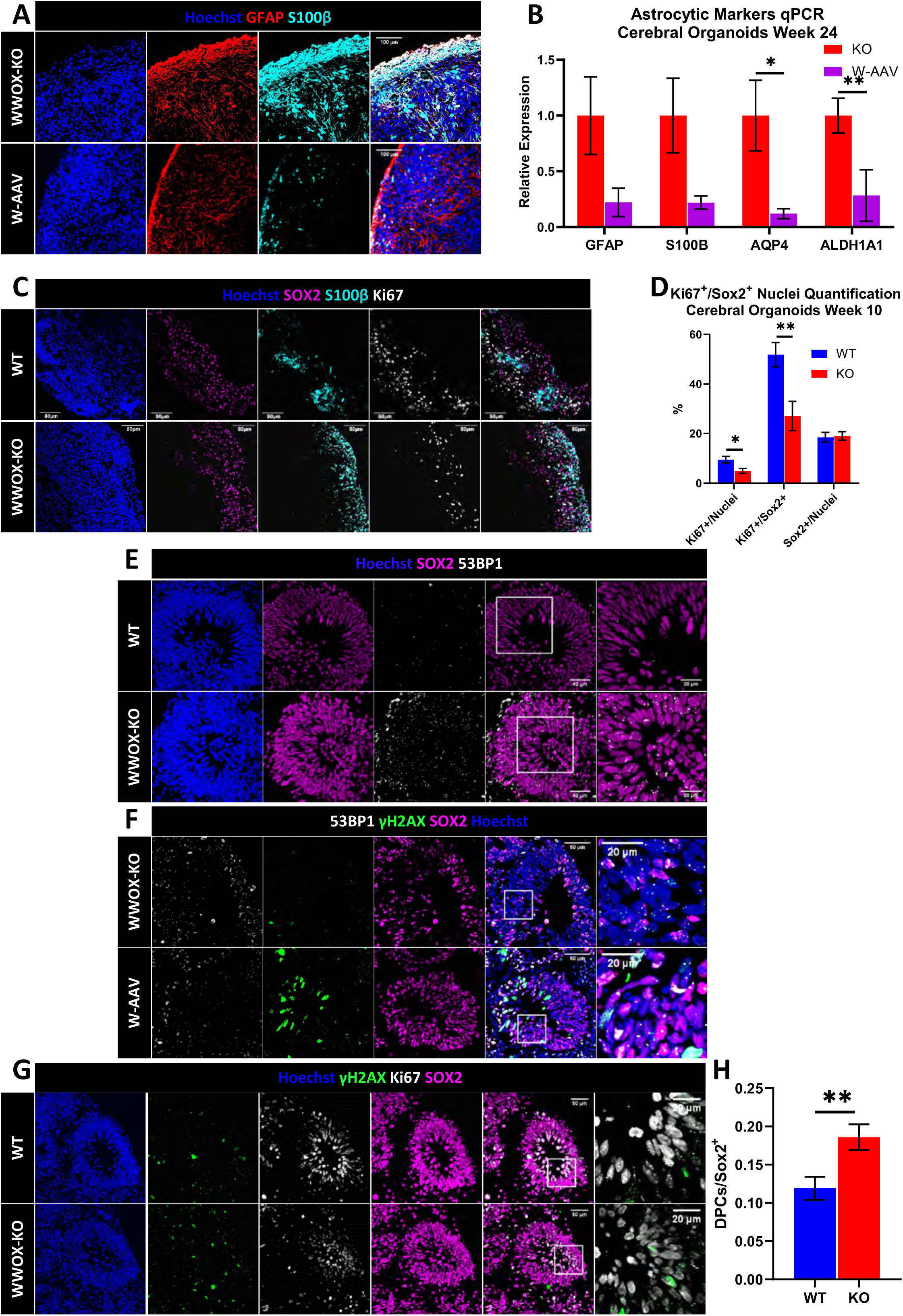
WWOX-KO Cerebral Organoids Showed Impaired Astrogenesis and DNA-damage Response. A) IF staining of the astrocytic markers GFAP and S100β in week 24 W-AAV COs compared to the age matched WWOX-KO COs *(W-AAV: n=5, KO: n=2)*. B) qPCR quantifying astrocytic markers in week 24 COs *(W-AAV: n=3, KO: n=3).* Data are represented as mean ± SEM. C) IF staining of week 10 COs for the proliferation marker Ki67 localized with S100β *(WT: n=4, KO: n=3)*. D) Quantification of (A). Data are represented as mean ± SEM. E) Staining for the DNA damage marker p53-Binding protein 1 (53BP1) in week 6 COs in the nuclei of cells in the VZ at physiological conditions *(WT: n=8 organoids from 3 individual batches, KO: n=12 organoids from 3 individual batches)*. F) Week 6 COs stained for the DNA damage markers γH2AX and 53BP1. Comparison of the foci demonstrated less sites of damage in the nuclei of cells in the VZ of W-AAV COs at physiological conditions *(W-AAV: n=4, KO: n=3)*. G) Week 10 COs stained for the Ki67 and γH2AX *(WT: n=4, KO: n=3)*. H) (H) Quantification of γH2AX+/Ki67+ double positive cells (DPCs) normalized to Sox2+ cells, corresponding to the size of the VZ, seen in (G).

**Figure S4.**
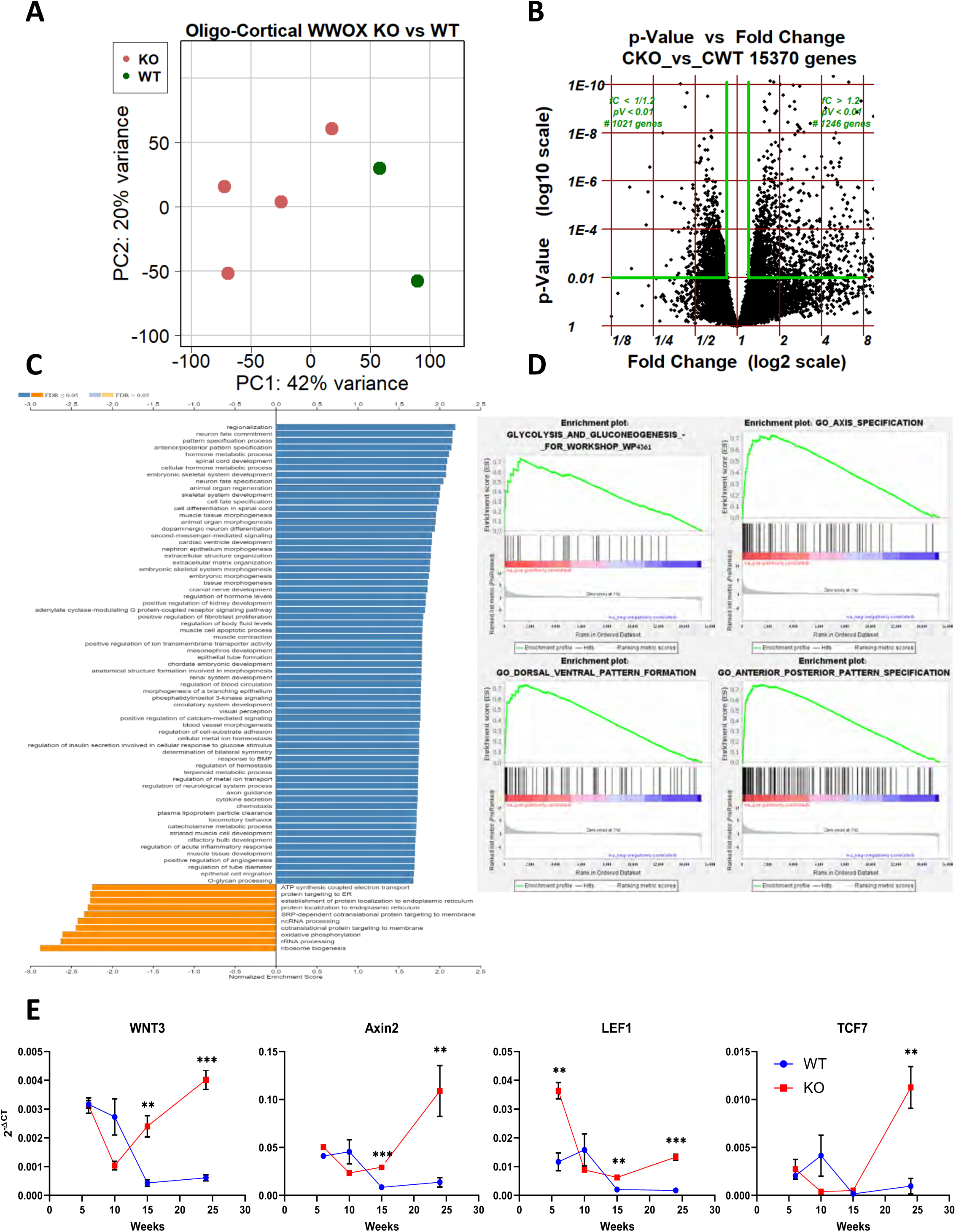
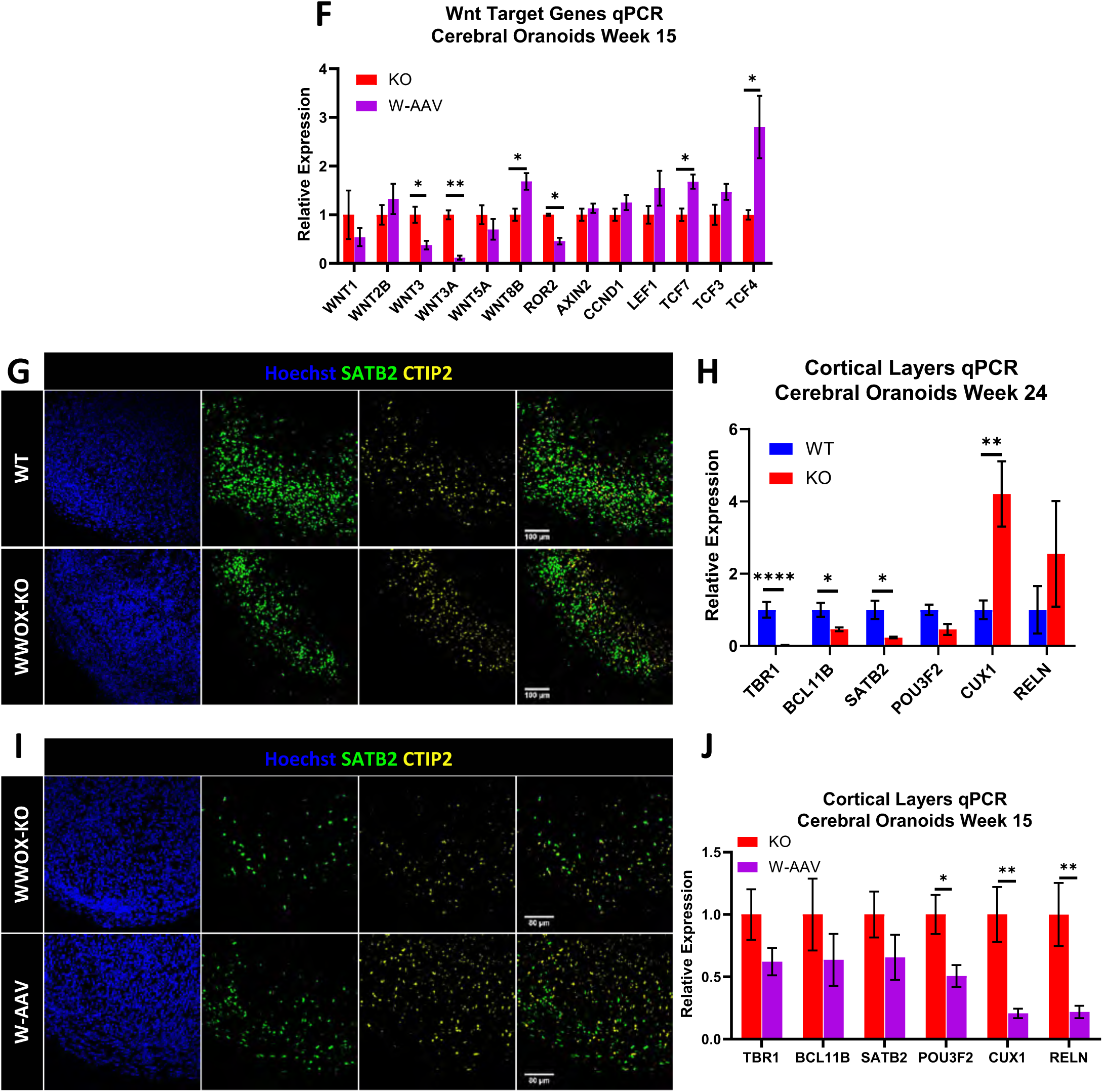
Cerebral Organoids RNA-sequencing Revealed Major Differentiation Defects. A) Principal component analysis (PCA) plot using the top two principal components, showing the RNA-seq data form week 15 COs. PCA revealed two distinct clusters corresponding to the biological identity of the sample – WT or WWOX-KO. B) Volcano plot representing differentially expressed genes in the sequenced COs based on fold change (x-axis) and p-value (y-axis). C) Gene ontology (GO) enrichment analysis for the top 3,000 differentially expressed genes. D) Gene-set enrichment analysis (GSEA) further emphasized activation of processes related to axis specification and glycolysis and gluconeogenesis. E) qPCR analysis of the kinetics of Wnt-related genes at week 6, 10, 15 and 24 COs. (WT W6: n=3, KO W6: n=3, WT W10: n=3, KO W10: n=3, WT W15: n=4, KO W15: n=4, WT W24: n=4, KO W24: n=3) F) qPCR quantifying the transcript levels of genes related to the Wnt signaling pathway in week 15 COs, showing downregulation of some genes. Data are represented as mean ± SEM. G) Immunofluorescent staining of cortical layers markers SATB2 and CTIP2 in week 24 COs *(WT: n=4, KO: n=2)*. H) qPCR measurement of the expression of human cortical layers markers *(WT: n=4* from 2 individual batches*, KO: n=3).* Data are represented as mean ± SEM. I) Week 12 WWOX-KO and W-AAV COs stained for the cortical layers’ markers SATB2 and CTIP2 (KO: n=3, W-AAV: n=3). J) qPCR quantifying the transcript levels of cortical layers markers. Data are represented as mean ± SEM.

**Figure S5.**
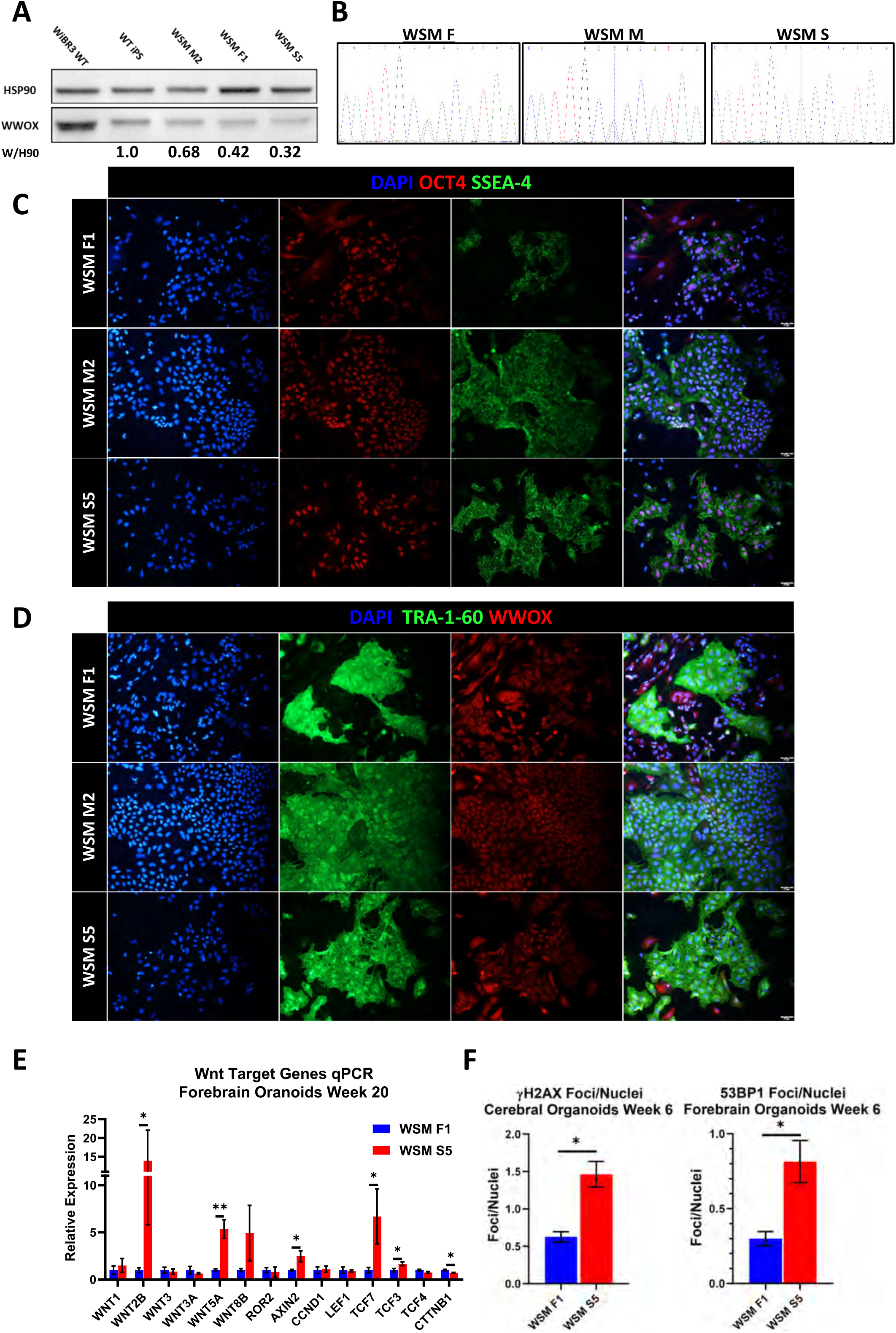
WWOX-Related Epileptic Encephalopathy Forebrain Organoids Presented Similar Phenotype to WWOX-KO COs. A) WB analysis of protein lysates from WT hESCs and iPSCs from a WOREE syndrome family [Mother (M), father (F) and son (S)] and an unrelated healthy donor (WT iPSC), showing the different levels of expression of WWOX, in correlation with the number of intact alleles in the genome. The numbers at the lower part of the number indicates quantification of the bands – WWOX is normalized to levels of HSP90 in the same line (W/H90), and fold change compared to the WT iPS line is indicated. B) Sanger sequencing of the c.517-2A>G mutation in the iPSCs shown in figure A. C) Expression of pluripotency genes (OCT4, SSEA-4) visualized using immunofluorescent antibodies validating the success of the reprogramming process in the iPSCs. D) Expression of the pluripotency gene TRA-1-60, together with WWOX, in iPSCs from the WOREE syndrome Family. E) Expression level of Wnt pathway related genes at mRNA levels quantified using qPCR in week 20 FOs *(WSM F1: n=4, WSM S5: n=3).* Data are represented as mean ± SEM. F) Quantification of γH2AX foci in the nuclei of cells composing the innermost layer of the VZ, normalized to the total number of nuclei in this layer. Data are represented as mean ± SEM.

**Figure S6.**
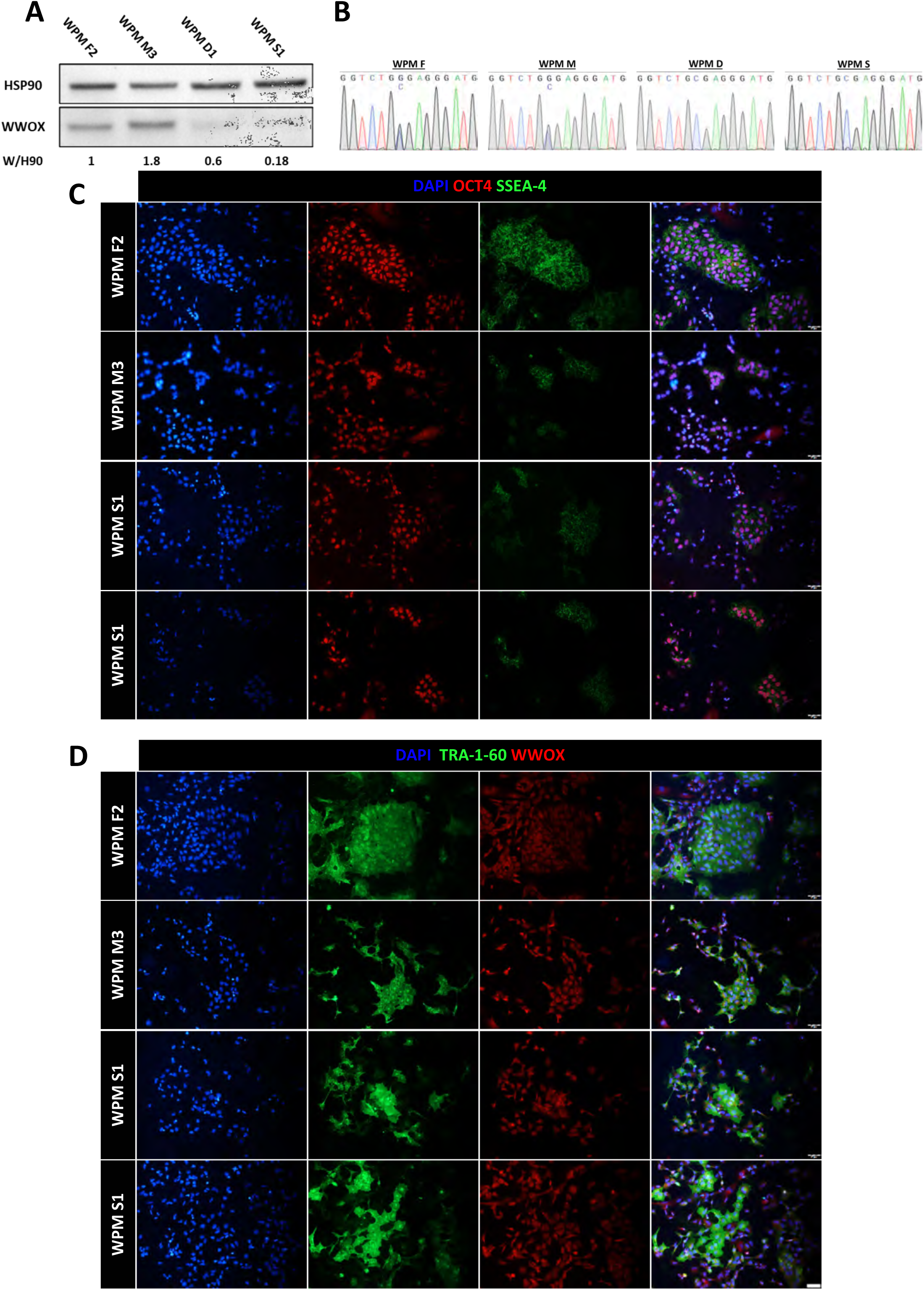
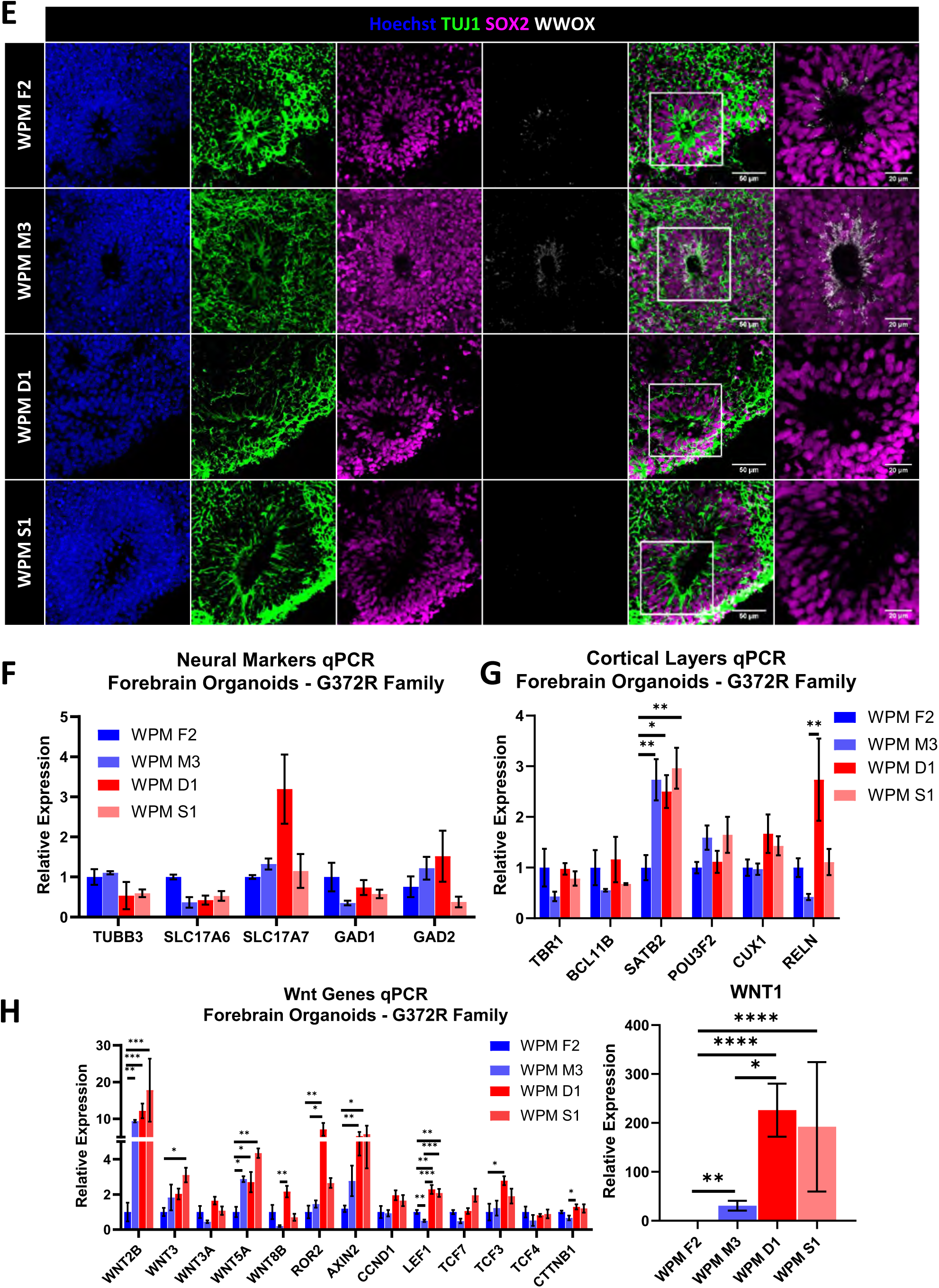
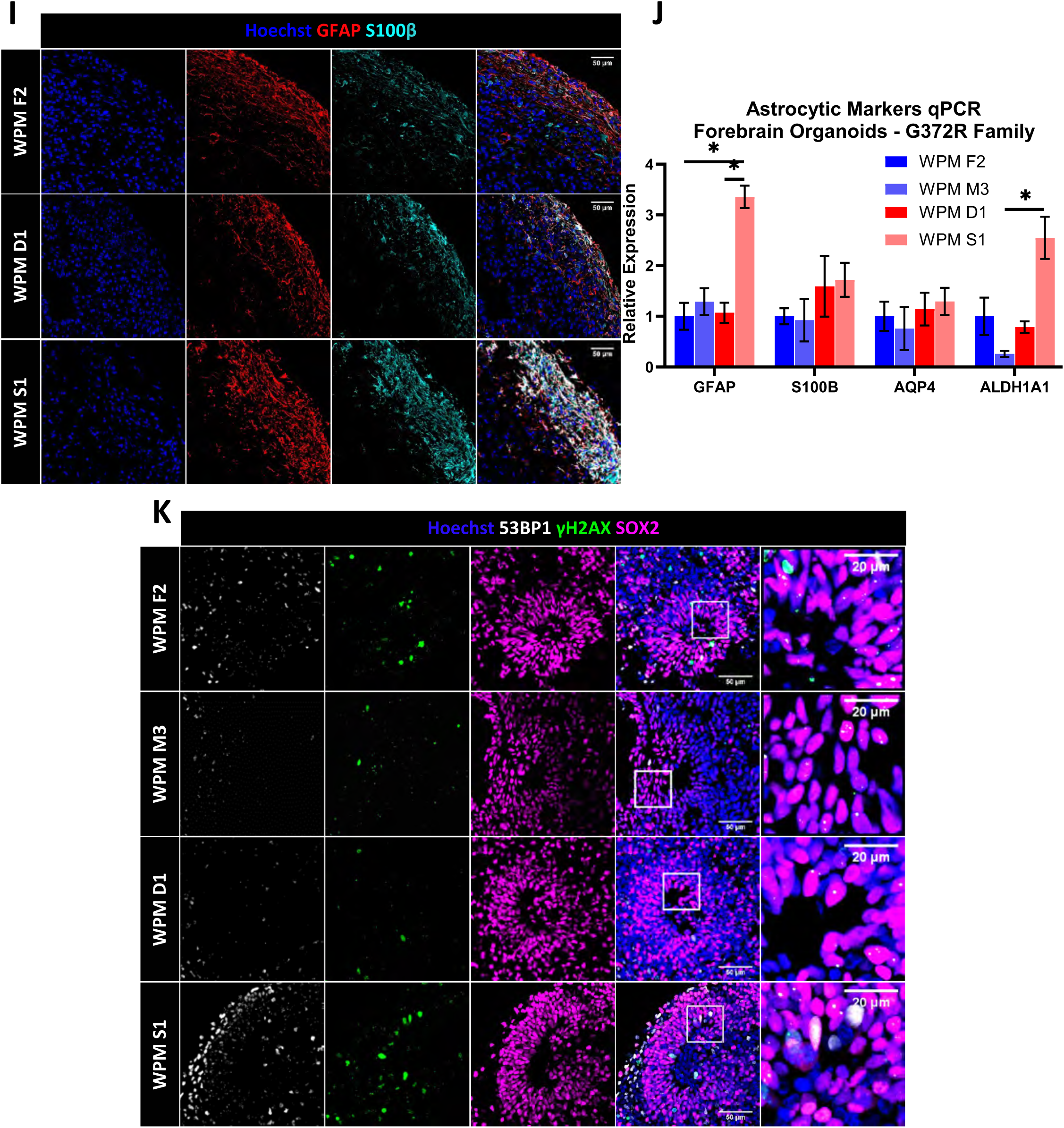
Spinocerebellar Ataxia Type 12 Forebrain Organoids Presented a Milder Phenotype Compared to WOREE FOs. PBMCs were isolated from SCAR12 (G372R mutation) patients and from their healthy parents, were reprogrammed into iPSCs, and subsequently were differentiated into forebrain organoids. A) Immunoblot of protein lysates from SCAR12 family (Mother (M), father (F), daughter (D) and son (S)) showing the reduced levels of WWOX, in correlation with the number of intact alleles in the genome. The numbers at the lower part of the number indicates quantification of the bands – WWOX is normalized to levels of HSP90 in the same line (W/H90), and fold change compared to WPM F2 line is indicated. B) Sanger sequencing of the c.1114G>C mutation in the iPSCs shown in figure A. C) Expression of pluripotency genes (OCT4, SSEA-4) visualized using immunofluorescent antibodies validating the success of the reprogramming process in the iPSCs. D) Expression of the pluripotency gene TRA-1-60, together with WWOX, in iPSCs from the SCAR12 syndrome Family. E) Week 6 forebrain organoids (FOs) of the healthy, heterozygote father (WPM F2) and mother (WPM M3), and their sick homozygotes daughter (WPM D1) and son (WPM S1) stained for WWOX expression. F) qPCR for the assessment of the expression of different neural markers in week 20 FOs (WPM F2: *n=4, WPM M3: n=3 WPM D1: n=4, WPM S1: n=3)*. G) qPCR for the measurement of expression levels of cortical layers markers in 20 weeks FOs (WPM F2: *n=4, WPM M3: n=3 WPM D1: n=4, WPM S1: n=3*). H) Expression level of Wnt pathway related genes at mRNA levels quantified using qPCR in week 20 FOs (WPM F2: *n=4, WPM M3: n=3 WPM D1: n=4, WPM S1: n=3*). F-H: Data are represented as mean ± SEM. I) Week 20 FOs stained for the astrocytic markers GFAP and S100β, showing comparable levels of expression in week 20 WPM D1 and WPM S1 FOs compared to the age matched WPM F2 FOs (WPM F2: *n=3, WPM D1: n=3, WPM S1: n=3)*. J) qPCR quantifying the transcript levels of astrocytic markers in week 20 FOs (WPM F2: *n=4, WPM M3: n=3 WPM D1: n=4, WPM S1: n=3).* Data are represented as mean ± SEM. K) Week 6 FOs stained for the DNA damage markers γH2AX and 53BP1. Comparison of the foci demonstrated no significant difference in sites of damage in the nuclei of cells in the VZ at physiological conditions *(WPM F2: n=1, WPM M3: n=2, WPM D1 n=2, WPM S1: n=2)*.

## Notes

### Competing Interest Statement

The authors have declared no competing interest.

