## Supplemental tables 3-5 for "Modeling Genetic Epileptic Encephalopathies using Brain Organoids"

**Supplementary Table 1 – 1,246 RNA-seq Upregulated Genes**

See Attached Supplementary excel file.

**Supplementary Table 2 – 1,021 RNA-seq Downregulated Genes**

See Attached Supplementary excel file.

**Supplementary Table 3 – sgRNA Sequences**

| **Name** | **Sequence 5'—>3'** | **Location** | **Reference** |
| --- | --- | --- | --- |
| WWOX sgRNA | CACCGCATGGCAGCGCTGCGCTACG | Exon1 | Abdeen et al., 2018 |
| AAVS1 sgRNA | GTCACCAATCCTGTCCCTAG | - | Guernet *et al.*, 2016 |

**Supplementary Table 4 – List of Antibodies Used in the Study**

| **Antibody** | **Host** | **Catalog number** | **Dilution** | **Application** | **Notes** |
| --- | --- | --- | --- | --- | --- |
| Cleaved Caspase 3 (D174) | Rabbit | 9661S | 1:300 | IF |  |
| 53BP1 (H300) | Rabbit | sc-22760 | 1:200 | IF |  |
| gH2AX (S139) | Rabbit | 9720S | 1:200 | IF |  |
| gH2AX (S139) | Mouse | ab26350 | 1:1000 | IF |  |
| Tuj1 | Mouse | 81202 | 1:1000 | IF |  |
| NeuN | Mouse | MAB377 | 1:400 | IF |  |
| GAD67 | Mouse | MAB5406 | 1:1000 | IF |  |
| VGLUT1 | Rabbit | ABN1647 | 1:500 | IF |  |
| CTIP2 | Rat | ab18465 | 1:300 | IF |  |
| SATB2 | Mouse | ab51502 | 1:300 | IF |  |
| GFAP | Mouse | MAB360 | 1:500 | IF |  |
| S100B | Rabbit | ab52642 | 1:200 | IF |  |
| Ki67 | Rabbit | MA5-14520 | 1:200 | IF |  |
| Sox2 | Rat | 14-9811-80 | 1:1000 | IF |  |
| TBR2 | Rabbit | ab2283 | 1:300 | IF |  |
| SSEA4 | Mouse | ab16287 | 1:100 | IF |  |
| TRA-1-60 | Mouse | ab16288 | 1:500 | IF |  |
| OCT3/4 | Rabbit | sc-5279 | 1:400 | IF |  |
| Anti-mouse Alexa Fluor 488 | Goat | A11029 | 1:1000 | IF |  |
| Anti-rabbit Alexa fluor 568 | Goat | A11011 | 1:1000 | IF |  |
| Anti-rat Alexa Fluor 594 | Goat | A11007 | 1:1000 | IF |  |
| Anti-Rabbit Alexa Fluor 647 | Goat | A21244 | 1:1000 | IF |  |
| WWOX | Rabbit | See notes. | 1:10,000 | IF, WB | Aqeilan et al., 2004 |
| WWOX | Rabbit | HPA050992 | 1:500 | IF |  |
| HSP90 | Rabbit | 4874 | 1:1000 | WB |  |
| KAP-1 | Rabbit | A300-274A | 1:5000 | WB |  |
| B-catenin | Mouse | 610154 | 1:200 (IHC), 1:1000 WB | IHC, WB |  |
| GAPDH | Mouse | CB1001 | 1:10,000 | WB |  |

**Supplementary Table 5 – Primer Sequences**

| **Gene** | **Direction** | **Sequence** | **Marker** | **Product size** |
| --- | --- | --- | --- | --- |
| HPRT | F | TGACACTGGCAAAACAATGCA | Housekeeping gene | 94 |
|  | R | GGTCCTTTTCACCAGCAAGCT |  |  |
| UBC | F | ATTTGGGTCGCGGTTCTTG | Housekeeping gene | 133 |
|  | R | TGCCTTGACATTCTCGATGGT |  |  |
| SOX2 | F | TTCACATGTCCCAGCACTACCAGA | Progenitor cells | 80 |
|  | R | TCACATGTGTGAGAGGGGCAGTGTGC |  |  |
| PAX6 | F | ATTACTGTCCGAGGGGGTCT | Progenitor cells | 80 |
|  | R | TAGCCAGGTTGCGAAGAACT |  |  |
| TUBB3 (TUJ1) | F | ATTCTGGTGGACCTGGAAC | Neurons | 99 |
|  | R | CCCACTCTGACCAAAGATGA |  |  |
| VGLUT1 (SLC17A7) | F | ATCATGTCCACCACCAACG | Glutamatergic neurons | 86 |
|  | R | GAGTAGCCGACCACCAACAG |  |  |
| SLC17A6 (vGlut2) | F | TGGGGCTACATCATCACTCA | Glutamatergic neurons | 85 |
|  | R | GAAGTATGGCAGCTCCGAAA |  |  |
| GAD1 | F | TCAAGTTCTGGCTGATGTGG | GABAergic neurons | 75 |
|  | R | CCAGTTCCAGGCATTTGTTG |  |  |
| GAD2 | F | AGCTGCTCCAAAGTGGATGT | Inhibitory neurons | 99 |
|  | R | ATCTTGCAGAAACGCCAAAG |  |  |
| GFAP | F | AAGAGGAACATCGTGGTGAA | Astrocytes | 86 |
|  | R | CACATCACATCCTTGTGCTC |  |  |
| S100B | F | GCTGAAGAAATCCGAACTGAA | Astrocytes | 84 |
|  | R | TCCACAACCTCCTGCTCTTT |  |  |
| AQP4 | F | TACTGGTGCCAGCATGAATC | Astrocytes | 89 |
|  | R | GCCCAACCCAATATATCCAA |  |  |
| ALDH1A1 | F | CCGTGGCGTACTATGGATGC | Astrocytes | 82 |
|  | R | GCAGCAGACGATCTCTTTCGAT |  |  |
| RELN | F | CTCTCCTCCCGAGAACTCAT | Layer 1 | 96 |
|  | R | GGAATGAAGGTCACCACAAG |  |  |
| CUX1 | F | AAGCCGAAACCATAGCTCTT | Layer 2 | 80 |
|  | R | TGTCTCCTGCAGCTTTCTCT |  |  |
| POU3F2 (BRN2) | F | GAGATGGCAAGCACTGAGAT | Layer 3 | 120 |
|  | R | TGTGCCAACTCTAAGCATCA |  |  |
| SATB2 | F | GTGTGTCCCAAGCTGTCTTT | Layer 4 | 87 |
|  | R | GAGGGTCTTCTTCCTTACGC |  |  |
| BCL11B (CTIP2) | F | GGCGATGCCAGAATAGATG | Layer 5 | 105 |
|  | R | CTCCACATGGTCAGCCTCT |  |  |
| TBR1 | F | GGCAGATGGTGGTTTTACAG | Layer 6 | 124 |
|  | R | TCAGGGAAAGTGAACGTCTG |  |  |
| WNT1 | F | CACCTCTTCGGCAAGATCG | Wnt Member | 116 |
|  | R | TCGATGGAACCTTCTGAGCA |  |  |
| WNT2B | F | ATCCACTACGGTGTCCGTTT | Wnt Member | 105 |
|  | R | GCGACCACAGCGGTTATTAT |  |  |
| WNT3A | F | ACATCGAGTTTGGTGGGATG | Wnt Member | 184 |
|  | R | CACCAGCATGTCTTCACCTC |  |  |
| WNT3 | F | ACTTCGGCGTGTTAGTGTCC | Wnt Member | 175 |
|  | R | CAGCAGGTCTTCACCTCACA |  |  |
| WNT5A | F | GCCCAGGTTGTAATTGAAGC | Wnt Member | 173 |
|  | R | CCGATGTACTGCATGTGGTC |  |  |
| WNT8B | F | TTTACTCCAGCAGTGTGGCA | Wnt Member | 144 |
|  | R | CTGTCTCCCGATTGGCACT |  |  |
| ROR2 | F | GGGGGAGATTGAAAACCGAA | Wnt receptor (Non canonical) | 112 |
|  | R | AAACACGAAGTGGCAGAAGG |  |  |
| Axin2 | F | TACACTCCTTATTGGGCGATCA | Wnt target | 151 |
|  | R | TTGGCTACTCGTAAAGTTTTGGT |  |  |
| CCND1 | F | GCTGCGAAGTGGAAACCATC | Wnt target | 135 |
|  | R | CCTCCTTCTGCACACATTTGAA |  |  |
| LEF1 (TCF7L3) | F | CCGTGAAGAGCAGGCTAAAT | Wnt target | 156 |
|  | R | CTTGGACCTGTACCTGATGC |  |  |
| TCF7 (TCF-1) | F | ATCAGCCAGAAGCAAGTTCA | Wnt target | 131 |
|  | R | CCTAGCATCAAGGATGGGTG |  |  |
| TCF7L1 (TCF-3) | F | AACGAGTCGGAGAACCAGAG | Wnt target | 151 |
|  | R | CAGGGTACGGGGGTCCTTTA |  |  |
| TCF7L2 (TCF-4) | F | CGTAGACCCCAAAACAGGAA | Wnt target | 171 |
|  | R | TAGGGGTGTCTGAATCCTCC |  |  |
| CTNNB1 | F | CTGAGGACAAGCCACAAGAT | Wnt target | 122 |
|  | R | CTGGGCACCAATATCAAGTC |  |  |
